# Aphid resistance segregates independently of cardiac glycoside and glucosinolate content in an *Erysimum cheiranthoides* (wormseed wallflower) F2 population

**DOI:** 10.1101/2024.01.11.575310

**Authors:** Mahdieh Mirzaei, Gordon C. Younkin, Adrian F. Powell, Martin L. Alani, Susan R. Strickler, Georg Jander

## Abstract

Plants in the genus *Erysimum* produce both glucosinolates and cardiac glycosides as defense against herbivory. Two natural isolates of *Erysimum cheiranthoides* (wormseed wallflower) differed in their glucosinolate content, cardiac glycoside content, and resistance to *Myzus persicae* (green peach aphid), a broad generalist herbivore. Both classes of defensive metabolites were produced constitutively and were not induced further by aphid feeding. To investigate the relative importance of glucosinolates and cardiac glycosides in *E. cheiranthoides* defense, we generated an improved genome assembly, genetic map, and segregating F2 population. Genotypic and phenotypic analysis of the F2 plants identified quantitative trait loci affecting glucosinolates and cardiac glycosides, but not aphid resistance. The abundance of most glucosinolates and cardiac glycosides was positively correlated in the F2 population, indicating that similar processes regulate their biosynthesis and accumulation. Aphid reproduction was positively correlated with glucosinolate content. Although overall cardiac glycoside content had little effect on aphid growth and survival, there was a negative correlation between aphid reproduction and helveticoside abundance. However, this variation in defensive metabolites could not explain the differences in aphid growth on the two parental lines, suggesting that processes other than the abundance of glucosinolates and cardiac glycosides have a predominant effect on aphid resistance in *E. cheiranthoides*.

## 1. Introduction

Most plants in the Brassicaceae rely on glucosinolates as their primary chemical defense against insect herbivory. These specialized metabolites are stored as inactive glucosides and are cleaved by myrosinases (thioglucosidases) during insect feeding to produce toxic and deterrent breakdown products [1]. In the 90 million years since the evolution of glucosinolate biosynthesis in the Brassicaceae [2], several crucifer-feeding specialist herbivores, including *Plutella xylostella* (diamondback moth), *Pieris rapae* (white cabbage butterfly), *Brevicoryne brassicae* (cabbage aphid), *Phyllotreta striolata* (striped flea beetle), and *Athalia rosae* (turnip sawfly), have evolved mechanisms to avoid or detoxify these plant defenses [3–9]. Broad generalist herbivores, such as *Trichoplusia ni* (cabbage looper) and *Myzus persicae* (green peach aphid), also feed readily on glucosinolate-containing plants [10–12].

Some Brassicaceae produce not only glucosinolates, but also additional chemical defenses that provide protection against specialist herbivores that are resistant to glucosinolates. One example of this more recent evolution of a second chemical defense is the accumulation of cardiac glycosides in the genus *Erysimum* [13–17]. Cardiac glycosides, a diverse group of metabolites that act as allosteric inhibitors of essential Na^+^/K^+^-ATPases in animal cells, are characteristic and well-studied herbivore defenses in *Digitalis* spp. (Plantaginaceae; foxglove) [18] and *Asclepias* spp. (Apocynaceae; milkweed) [19]. Within the Brassicaceae, cardiac glycosides have been found almost exclusively in the *Erysimum* genus [13,14,20]. Phylogenetic studies involving more than 100 *Erysimum* species suggest rapid speciation in this genus after the evolution of cardiac glycoside biosynthesis about three million years ago [21–23].

An available genome sequence, transcriptomes, and metabolomic data for *Erysimum cheiranthoides* (wormseed wallflower) [24] facilitate use of this species for studying the combined function of cardiac glycosides and glucosinolates in plant defense. The most abundant cardiac glycosides in *E. cheiranthoides* are mono- and diglycosides of digitoxigenin, cannogenol, cannogenin, and strophanthidin [13,24]. Glucosinolates with side chains derived from tryptophan and methionine, which are abundant in the genetic model plant *Arabidopsis thaliana* (Arabidopsis), are also present in *E. cheiranthoides* [25,26]. Although analysis of the *E. cheiranthoides* genome sequence identified homologs of most Arabidopsis glucosinolate biosynthesis genes [23] their specific functions have not been investigated.

Unlike in the case of milkweeds, which have a community of highly adapted herbivores that are largely impervious to inhibition by cardiac glycosides [27], there are no known *Erysimum-*specialist herbivores that are resistant to both glucosinolates and cardiac glycosides. The relatively recent evolution of cardiac glycoside production in *Erysimum* may account for the absence of such specialized herbivores. Experiments with two crucifer-specialist lepidopterans, *P. rapae* and *Pieris napi* (green-veined white butterfly), showed that *Erysimum* cardiac glycosides deter both oviposition and feeding [15,28–34]. However, larvae of *P. xylostella*, another crucifer-specialist lepidopteran, have been reported on *E. cheiranthoides* in field experiments [35].

*Myzus persicae*, a broad generalist herbivore, is able to feed on *E. cheiranthoides* in the laboratory and in nature [15,35,36], indicating that this species has some tolerance for both glucosinolates and cardiac glycosides. When feeding on Arabidopsis, methionine-derived aliphatic glucosinolates pass through *M. persicae* largely intact, whereas tryptophan-derived indole glucosinolates are activated in the aphid gut [37]. Although Arabidopsis *cyp79B2 cyp79B3* mutants, which lack indole glucosinolates, are more sensitive to *M. persicae* [38], *tgg1 tgg2* mutants, which are deficient in glucosinolate-activating myrosinases, are not [39].

In the current study, we conducted experiments with two *E. cheiranthoides* accessions, Elbtalaue and Konstanz, with differing glucosinolate content, cardiac glycoside content, and aphid resistance. By measuring these traits in a segregating Elbtalaue × Konstanz F2 population, we mapped genetic loci affecting the abundance of glucosinolates and cardiac glycosides. Additionally, we used this mapping population to investigate the relative importance of these compounds in *E. cheiranthoides* defense against *M. persicae* feeding.

## 2. Results

### 2.1 Phenotypic differences between the Elbtalaue and Konstanz accessions

We investigated two inbred *E. cheiranthoides* accessions, Elbtalaue and Konstanz, for variation in aphid resistance, glucosinolate accumulation, and cardiac glycoside accumulation. In no-choice assays, *M. persicae* survival and reproduction were higher on Konstanz than on Elbtalaue (Figure 1A,B). Similarly, aphids showed a preference for detached leaves of Konstanz plants relative to those of Elbtalaue in choice assays (Figure 1C). We measured the relative abundance of glucosinolates (Figure 2) and cardiac glycosides (Figure 3) in the Elbtalaue and Konstanz accessions, as well as in aphids feeding on the leaves of these plants. Among the 8 glucosinolates and 7 cardiac glycosides that we reliably detected in *E. cheiranthoides* leaves, none had significantly increased abundance after 24 hours of aphid feeding. However, some glucosinolates [4-hydroxyindol-3-ylmethylglucosinolate (4HI3M), 3-methylsulfonylpropylglucosinolat (3-MSOP), and 4-methylsulfonylbutylglucosinolate (4MSOB)] exhibited a transient increase in abundance after 1 h and then decreased to background levels after 24 h (Figure 2C, I, K).

**Figure 1.**
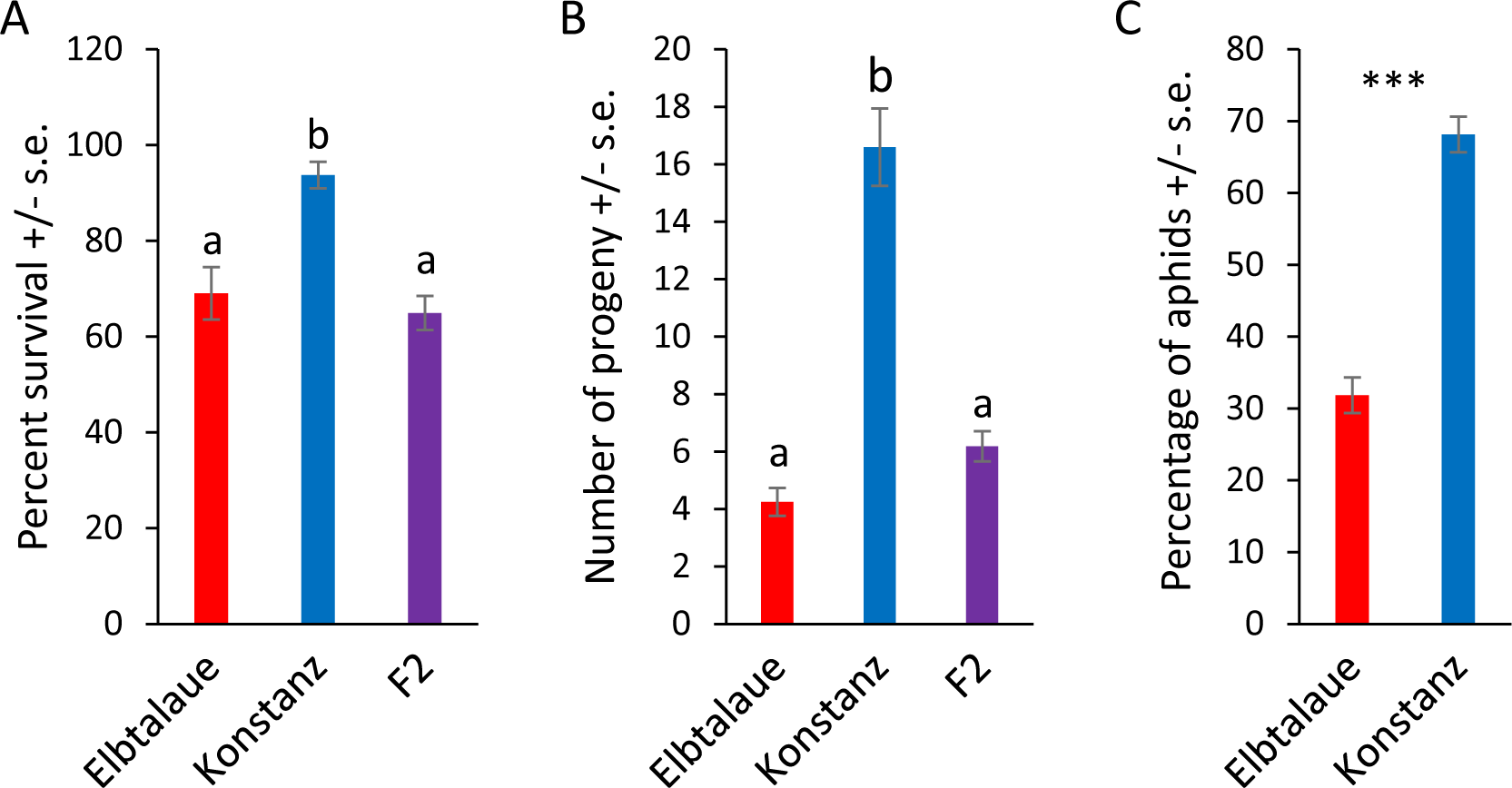
Aphid performance on two *Erysimum cheiranthoides* accessions. (A) Survival and (B) reproduction of *Myzus persicae* on *E. cheiranthoides* accessions Elbtalaue (N = 56), Konstanz (N = 32), and an F2 population (N = 155). Three nymphs were placed in each cage and the number of surviving adults and progeny produced were counted after 10 days. Mean +/− s.e., different letters indicate P < 0.05, ANOVA followed by Tukey’s HSD test. (C) Choice assays with detached leaves in Petri dishes. Mean +/− s.e. of N = 30. ***P < 0.005, paired *t*-test.

**Figure 2.**
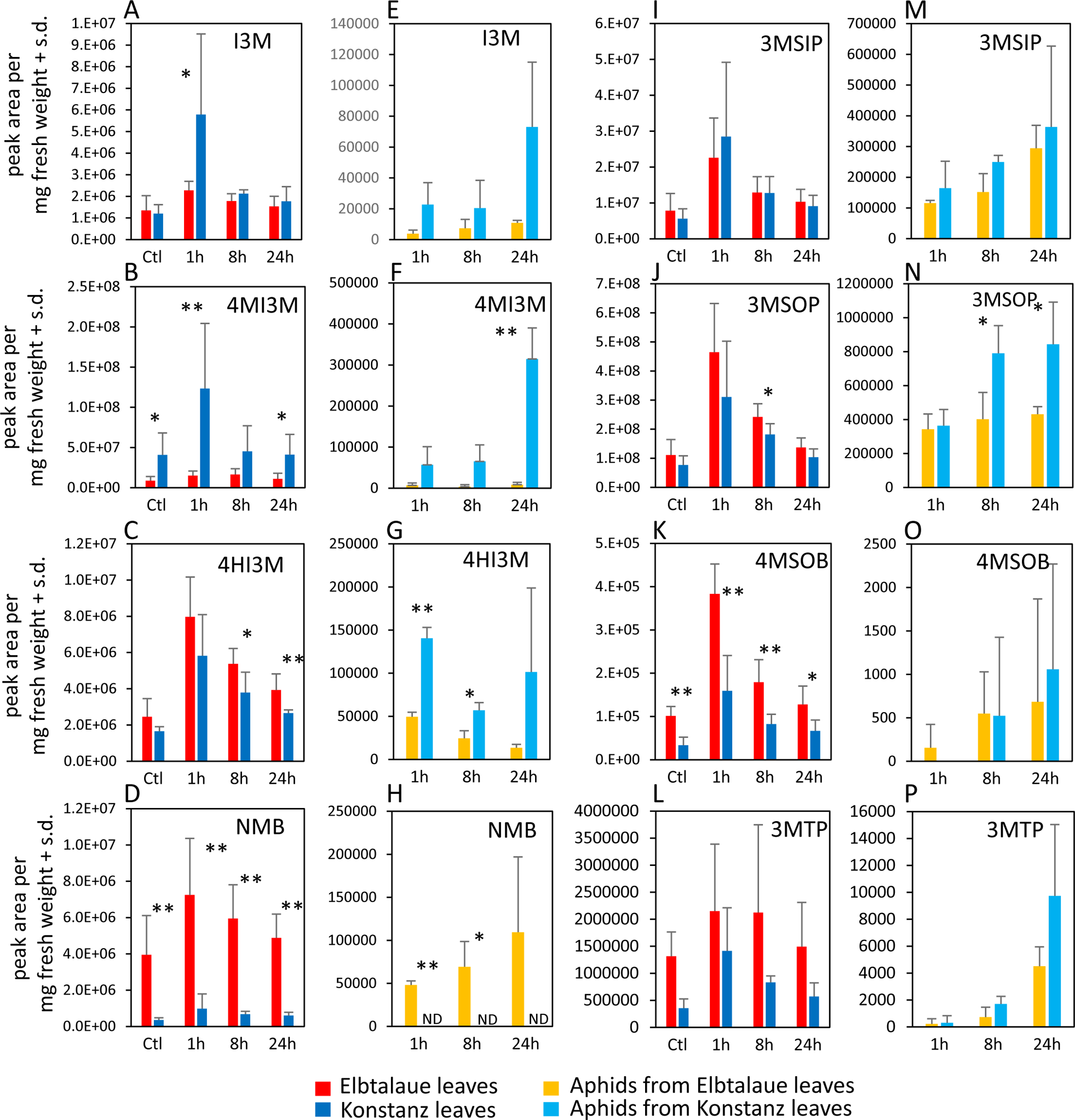
Glucosinolate content of two *Erysimum cheiranthoides* accessions, Elbtalaue and Konstanz, and aphids feeding on these plants. Samples were collected from uninfested control plants (Ctl) and after 1, 8, and 24 h of *Myzus persicae* feeding. (A-C) Indole glucosinolates (D, I-L) aliphatic glucosinolates in plant samples. (E-G) Indole glucosinolates (H, M-P) aliphatic glucosinolates in aphid samples. Mean ± s.d. of N = 6 (plant samples) or 3 (aphid samples), *P < 0.05, **P < 0.01, *t-*test comparing Elbtalaue and Konstanz samples. ND = not detected. Glucosinolate side chain abbreviations: I3M = indol-3-ylmethyl, 4HI3M = 4-hydroxyindol-3-ylmethyl, 4MI3M = 4-methoxyindol-3-ylmethyl, 3MSIP = 3-methylsulfinylpropyl, 3MSOP = 3-methylsulfonylpropyl, 4MSOB = 4-methylsulfonylbutyl, 3MTP = 3-methylthiopropyl, and NMB = n-methylbutyl.

**Figure 3.**
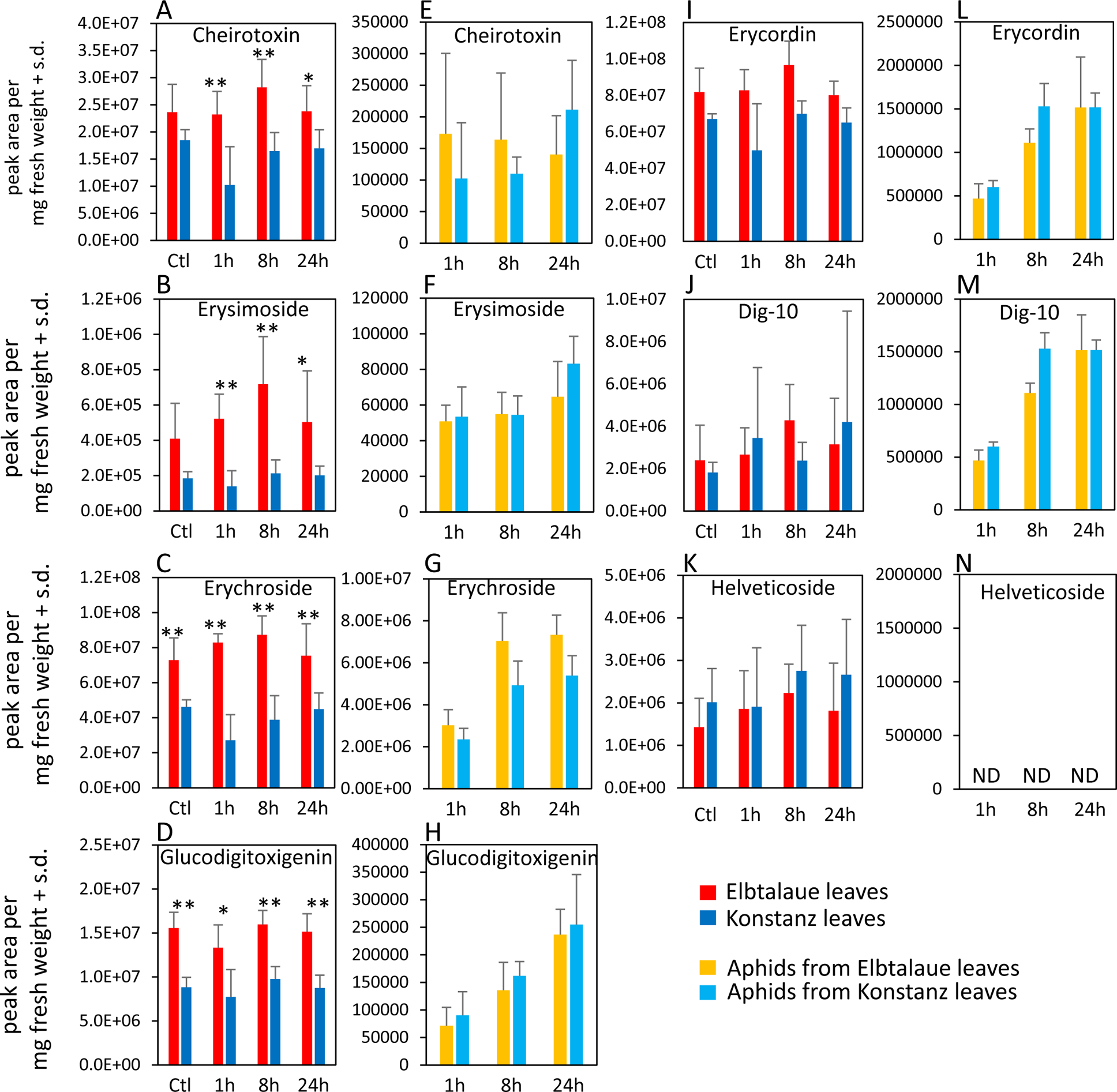
Cardiac glycoside content of two *Erysimum cheiranthoides* accessions, Elbtalaue and Konstanz, and aphids feeding on these plants. Samples were collected from uninfested control plants (Ctl) and after 1, 8, and 24 h of *Myzus persicae* feeding. (A-D, I-K) cardiac glycosides in plant samples, (E-H, L-N) cardiac glycosides in aphid samples. Helveticoside was not detected in aphid samples. Mean ± s.d. of N = 5-6 (plant samples) or 3 (aphid samples), *P < 0.05, **P < 0.01, *t-*test. comparing Elbtalaue and Konstanz samples. ND = not detected.

Whereas 4-methoxyindol-3-ylmethylglucosinolate (4MI3M) was more abundant in Konstanz than in Elbtalaue, both constitutively and after aphid feeding, 4HI3M, was more abundant in Elbtalaue after 8 and 24 h of feeding (Figure 2B). The abundance of indole glucosinolates was generally higher in aphids on the Konstanz accession and, after 24 h feeding on Konstanz, the 4MI3M concentration was higher in aphids than at the 1 h and 4 h timepoints (P < 0.05, *t-*test; Figure 2F). 4MSOB was twice as abundant and n-methylbutylglucosinolate (NMB) was ten-fold more abundant in Elbtalaue than in Konstanz (Figure 2D). Likely due to the relatively low abundance of the aliphatic glucosinolate NMB in the Konstanz accession, this glucosinolate was not detected above background in assays of aphids collected from these plants (Figure 2H).

In the case of cardiac glycosides, cheirotoxin, erysimoside, erychroside, and glucodigitoxigenin, were all more abundant in Elbtalaue during the aphid feeding experiment (Figure 3A-D). However, there was no significant difference in the abundance of these cardiac glycosides in the bodies of aphids feeding from these plants (Figure 3E-H). Three additional cardiac glycosides, helveticoside, erycordin, and the structurally uncharacterized Dig-10 did not differ in abundance between Elbtalaue and Konstanz (Figure 3I-K). Uniquely among the detected cardiac glycosides, helveticoside was not detected by HPLC-MS in aphids that were feeding on either of the two *E. cheiranthoides* accessions (Figure 3N).

### 2.2 Correlation of aphid resistance with glucosinolate and cardiac glycoside content

To investigate genetic basis of variation in aphid resistance, glucosinolate content, and cardiac glycoside content, we generated an F2 population from a cross between the Elbtalaue and Konstanz accessions. Aphid survival and reproduction on F2 progeny were similar to those observed on Elbtalaue and significantly different from Konstanz (Figure 1A,B). This suggested that resistance was a dominant trait in this cross and that multiple loci contributed to the higher level of aphid resistance in Elbtalaue relative to Konstanz.

From the 155 F2 plants that we used for aphid bioassays (Figure 1), we subjected 83 to glucosinolate analysis, cardiac glycoside analysis, and transcriptome sequencing. After data normalization, we conducted Pearson correlation analysis to: 1) compare aphid resistance (progeny production) and metabolite content and 2) understand the correlation in the abundances of the different metabolites (Figure 4A). Most comparisons of cardiac glycoside and glucosinolate abundance showed a positive correlation. However, helveticoside abundance showed no significant correlation with the abundances of the measured glucosinolates. Aphid reproduction was significantly negatively correlated with the helveticoside abundance and positively correlated with glucosinolate abundance in the F2 population. There was no correlation between the abundance of the other measured cardiac glycosides with aphid reproduction. We confirmed the negative effect of helveticoside on aphid reproduction using an artificial diet assay (Figure 4B). The calculated IC50 concentration for aphid progeny production on artificial diet was 14 ng/µl, which is comparable to the helveticoside content of *E. cheiranthoides* leaves (∼20 ng/mg wet weight) [36].

**Figure 4.**
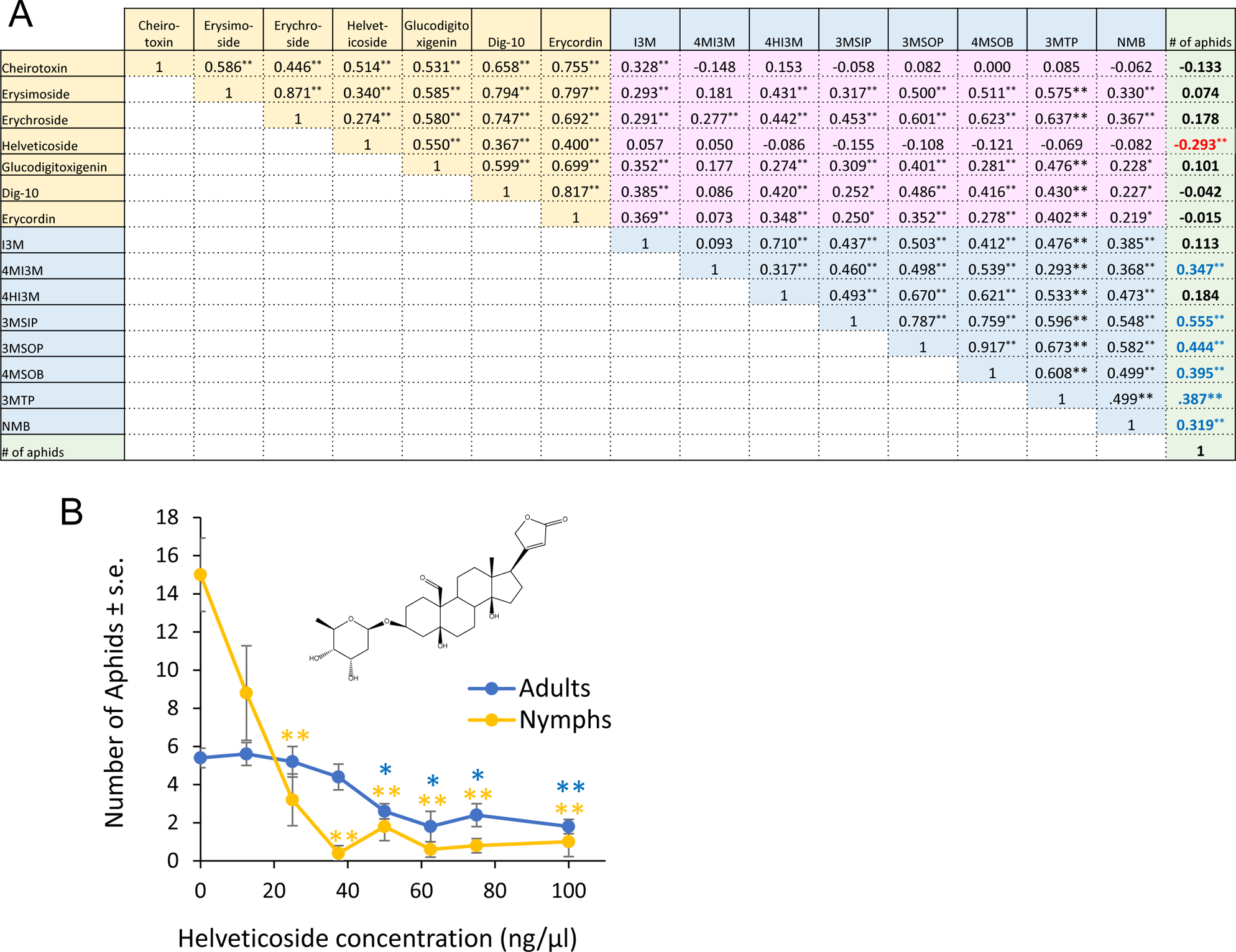
Correlation between glucosinolates, cardiac glycosides, and aphid reproduction. (A) Pearson correlation of cardiac glycoside content, glucosinolate content, and *Myzus persicae* progeny production on 83 F2 individuals from a cross between *Erysimum cheiranthoides* accessions Elbtalaue and Konstanz. The numbers in the boxes indicate correlation coefficients, with *P < 0.05, ** P < 0.01. Cardiac glycosides that are negatively correlated with aphid reproduction are indicated with red numbers, and glucosinolates that are positively correlated with aphid reproduction are indicted with blue numbers. Glucosinolate side chain abbreviations: I3M = indol-3-ylmethyl, 4HI3M = 4-hydroxyindol-3-ylmethyl, 4MI3M = 4-methoxyindol-3-ylmethyl, 3MSIP = 3-methylsulfinylpropyl, 3MSOP = 3-methylsulfonylpropyl, 4MSOB = 4-methylsulfonylbutyl, 3MTP = 3-methylthiopropyl, and NMB = n-methylbutyl. (B) *M. persicae* survival and reproduction on diet with helveticoside. Mean ± s.e. of N = 4. Mean ± s.e. of N = 5. *P < 0.05, **P < 0.005, Dunnett’s test relative to no-cardiac glycosides control for adults (blue) and nymphs (orange). Inset = chemical structure of helveticoside.

### 2.3 E. cheiranthoides genetic map

The previously published *E. cheiranthoides* genome (version 1.2, [24]) was constructed using 39.5 Gb of PacBio sequences and Hi-C proximity-guided assembly to orient 98.5% of the genome into eight scaffolds. We used transcriptome data from the F2 population to generate an *E. cheiranthoides* genetic map with 501 molecular markers (Figures S1, S2). With this genetic map, we re-scaffolded the assembled contigs for version 2.0 of the genome. A comparison of marker positions between versions 1.2 and 2.0 highlights several inversions and rearrangements, primarily on chromosomes 1, 4, 6, 7, and 8, that are corrected in the new genome assembly (Figure S3). Version 2.0 of the *E. cheiranthoides* genome has improved assembly statistics relative to previously published version 1.2 (Figure S4). In addition, we assembled 93 formerly unassigned contigs into a 154,508 bp chloroplast genome, which is similar to the 154,611 bp chloroplast genome described previously for a different isolate of *E. cheiranthoides* [40].

In some parts of the genome, the frequency of molecular markers is distorted from the expected 1:2:1 (Elbtalaue : Heterozygote : Konstanz) ratio for an F2 population (Figure S5). Particularly noteworthy is that Elbtalaue alleles are overrepresented across much of chromosome This segregation distortion could indicate that there is a selective advantage to specific parental alleles under our growth conditions. While conducting this research, we noticed that, relative to Elbtalaue, Konstanz seeds require longer cold stratification to achieve full germination. If loci affecting this trait are localized on chromosome 3 and the F2 population seeds was not cold-stratified long enough prior to planting, this could explain some of the unexpected allele frequencies in the F2 population.

### 2.4 Genetic mapping of defense traits

Using the newly assembled *E. cheiranthoides* genetic map (Figure S1) and 83 genotyped Elbtalaue x Konstanz F2 lines, we conducted quantitative trait locus (QTL) mapping of aphid survival, aphid progeny reproduction, cardiac glycoside abundance, and glucosinolate abundance. No significant QTL affecting aphid survival or progeny production on *E. cheiranthoides* F2 lines were identified. Significant genetic linkage was observed for only one cardiac glycoside, helveticoside (Figure 5A). The Konstanz allele of a locus on chromosome 8 causes an approximately two-fold increase in helveticoside abundance, an effect is likely recessive because F2 plants that are heterozygous at this locus have helveticoside levels similar to the Elbtalaue parent (Figure 5B). As there are no genes known to be involved specifically in helveticoside biosynthesis and the QTL mapping interval encompasses hundreds of genes, it is not yet possible to identify candidate loci affecting helveticoside accumulation.

**Figure 5.**
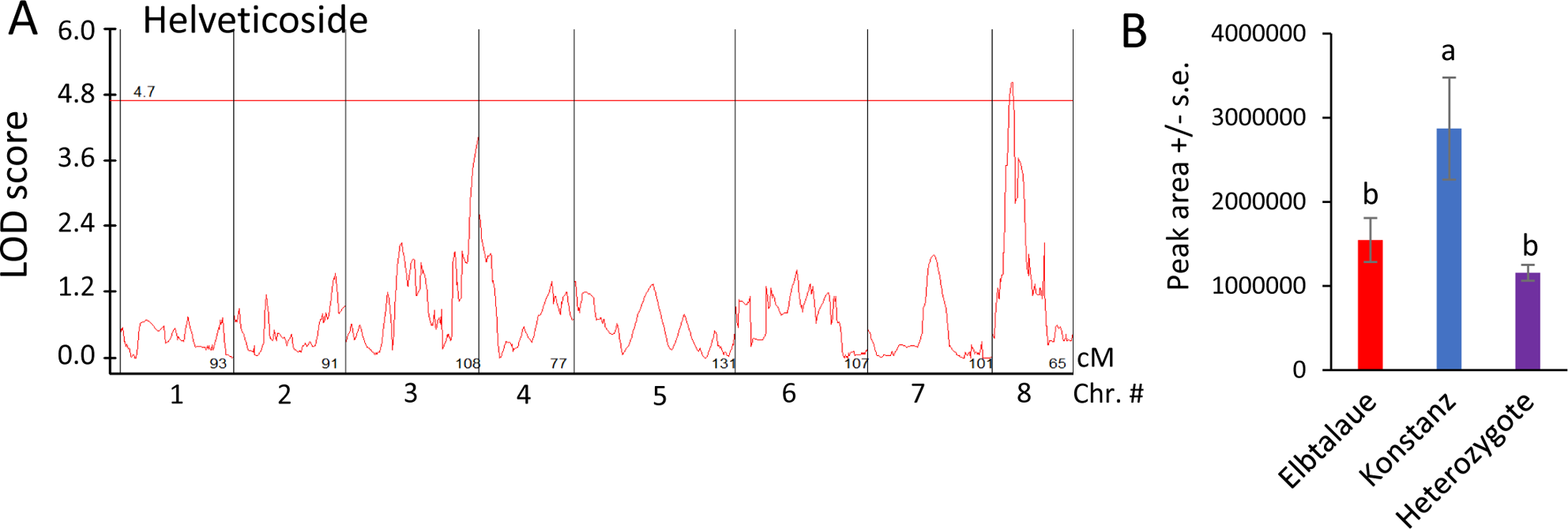
Quantitative trait locus (QTL) affecting helveticoside abundance in *Erysimum cheiranthoides*. (A) LOD plot of helveticoside abundance in an Elbtalaue x Konstanz F2 population (B) Helveticoside peak area at the chromosome 8 QTL, sorted by genotype. Mean +/− s.e. of N = 27 (Elbtalaue), 9 (Konstanz), and 45 (Heterozygote). Different letters indicates significant differences, P < 0.05, ANOVA followed by Tukey’s HSD test. The horizontal line in panel A is the 95% confidence level, calculated based on 500 permutations of the data.

NMB, the glucosinolate showing the greatest fold-difference between Elbtalaue and Konstanz (Figure 2D), has a significant QTL on chromosome 1, with a recessive allele in Elbtalaue causing increased foliar NMB accumulation (Figure 6A,B). Similar glucosinolates with five-carbon side chains derived from isoleucine have been described in *Boechera stricta* (Drummond’s rockcress) [41]. The relative incorporation of methionine and branched chain amino acids (valine and isoleucine) into glucosinolate side chains was associated with natural variation in CYP79F enzymes that catalyze the first step of the biosynthesis pathway. Analysis of *E. cheiranthoides* chromosome 1 in the area of the NMB QTL showed a *CYP79F* gene (Erche01g017900), with an encoded protein sequence that is similar to those from Arabidopsis, *B. stricta*, and *Brassica oleracea* (cabbage) (Figures S6A and S7). Erche01g017900 expression was not significantly different between the Ebltalaue and Konstanz accessions (P > 0.05; Figure S6B).

**Figure 6.**
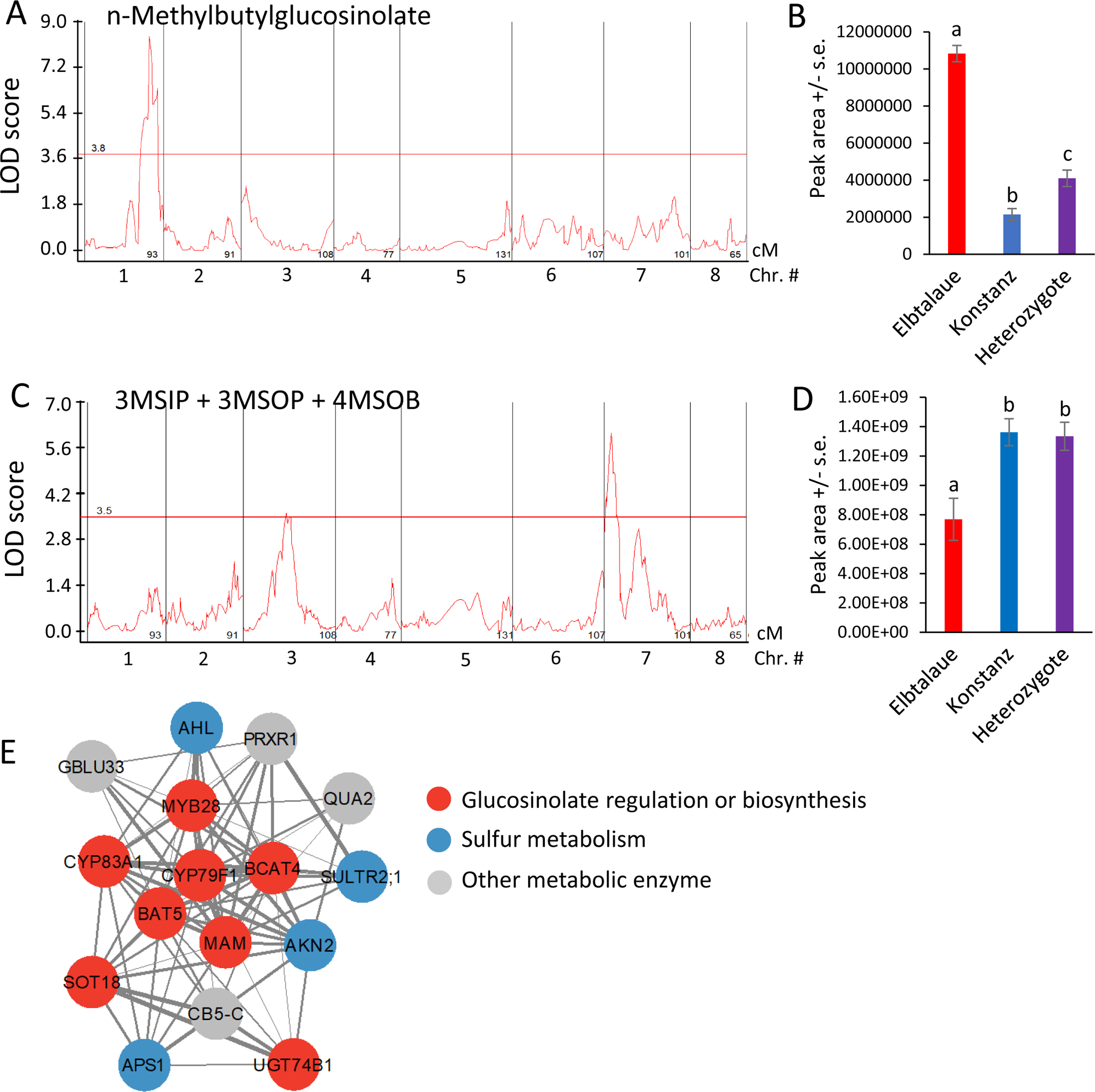
Quantitative trait loci (QTL) affecting *Erysimum cheiranthoides* aliphatic glucosinolate abundance. (A) LOD plot of n-methylbutylglucosinolate (NMB) abundance in an Elbtalaue x Konstanz F2 population (B) NMB peak area at the chromosome 1 QTL, sorted by genotype. Mean +/− s.e. of N = 15 (Elbtalaue), 21 (Konstanz), and 46 (Heterozygote). (C) LOD plot of 3-methylsulfinylpropylglucosinolate (3MSIP), 3-methylsulfonylpropylglucosinolate (3MSOP), and 4-methylsulfonylbutylglucosinolate (4MSOB) abundance in an Elbtalaue x Konstanz F2 population (D) Peak of the chromosome 7 quantitative trait locus (QTL), sorted by genotype. Mean +/− s.e. of N = 15 (Elbtalaue), 21 (Konstanz), and 46 (Heterozygote). Different letters indicates significant differences, P < 0.05, ANOVA followed by Tukey’s HSD test. Horizontal lines in panels A and C are 95% confidence levels, calculated based on 500 permutations of the data. (E) Co-expression network containing genes involved in glucosinolate biosynthesis. Figure was made using Cytoscape v3.9.1. Full gene descriptions are in Supplemental Table S1.

The predicted Erche01g017900 protein sequences from Elbtalaue and Konstanz differ at only one amino acid (Figure S6A). Whereas Elbtalaue has glycine at position 51, Konstanz has serine. In *B. stricta*, *Bs*BCMA1 and *Bs*BCMA3, the two CYP79F enzymes associated with branched-chain amino acid incorporation, have serine at this position, and *Bs*BCMA2, which preferentially catalyzes methionine incorporation, has glycine (Figure S6). At two other positions that have been associated with differential glucosinolate production in *B. stricta* [41], residues 135 and 536, the Konstanz and Elbtalaue proteins are identical and have the same amino acids as those found in *Bs*BCMA1 and *Bs*BCMA3 (Figure S6).

The accumulation of 4MSOB, 3-methylsulfinylpropylglucosinolate (3MSIP), and 3MSOP, which are predicted to be synthesized by a shared biosynthetic pathway [24], is highly correlated (Figure 4A). Although mapping the accumulation of each of these glucosinolates individually did not identify significant QTL at the 95% confidence level (Figure S8), the sum of these three glucosinolates had a significant QTL localized on chromosome 7 (Figure 6C), with the recessive Elbtalaue allele causing lower glucosinolate accumulation (Figure 6D).

We conducted mutual rank coexpression network analysis [42] to determine whether known homologs of known Arabidopsis glucosinolate biosynthesis genes are also co-expressed in *E. cheiranthoides.* This identified a network of co-expressed genes containing eight genes involved in aliphatic glucosinolate biosynthesis, four genes related to sulfur metabolism, and four additional genes encoding likely metabolic enzymes (Figure 6E and Supplemental Table S1). Several *E. cheiranthoides* genes encoding aliphatic glucosinolate biosynthetic genes have expression-level QTL between 3.0 and 3.4 Mbp on chromosome 8. Known Arabidopsis transcription factors regulating aliphatic glucosinolate biosynthesis include MYB28, MYB29, and MYB76 [43]. However, *E. cheiranthoides* homologs of these genes are not located in this part of the genome, suggesting that gene expression variation in our F2 population is regulated by some other mechanism that genetically maps to chromosome 7.

Indol-3-ylmethylglucosinolate (I3M) is hydroxylated to form 4HI3M and then methylated to form 4MI3M (Figure 7A). 4MI3M is significantly more abundant in the Konstanz parent than in the Elbtalaue parent of the F2 population. To determine whether there is genetic regulation of the relative 4MI3M content, we mapped the ratio of peak areas, (4MI3M)/(4MI3M + 4HI3M), as a quantitative trait (Figure 7B). For both of the detected QTL, the Konstanz allele caused higher relative 4MI3M accumulation (Figure 7C,D), with the Elbtalaue allele on chromosome 2 being recessive and the allele on chromosome 3 being dominant. The two Konstanz alleles had an additive effect on 4MI3M concentration (Figure 7E). To identify loci that influence indole glucosinolate hydroxylation, we mapped the ratio (4MI3M + 4HI3M)/(I3M + 4MI3M + 4HI3M) as a quantitative trait. This identified a dominant locus from the Konstanz genetic background on chromosome 7 that increased the relative abundance of modified indole glucosinolates (4MI3M + 4HI3M) (Figure 7F,G). The *E. cheiranthoides* homologs of Arabidopsis enzymes that catalyze I3M 4-hydroxylation [44,45], are encoded on chromosome 2 (Erche02g041710 and Erche02g041680). Therefore, cis-regulation or differences in enzymatic activity are unlikely to be the cause of this variation in the indole glucosinolate profile.

**Figure 7.**
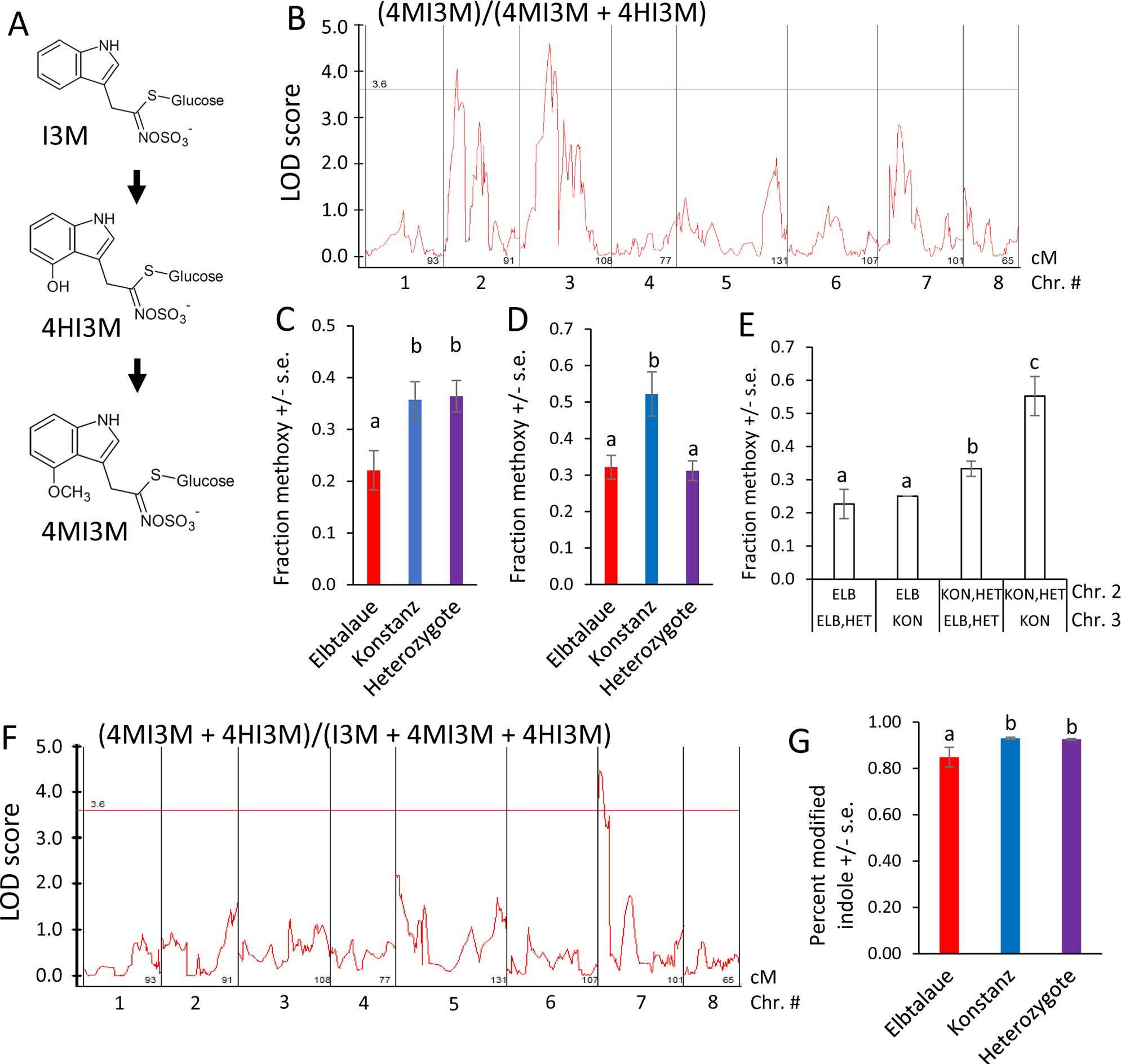
Quantitative trait loci (QTL) affecting *Erysimum cheiranthoides* indole glucosinolate abundance. (A) Pathway of indole glucosinolate modification by hydroxylation and O-methylation. Glucosinolate side chain abbreviations: I3M = indol-3-ylmethyl, 4HI3M = 4-hydroxyindol-3-ylmethyl, 4MI3M = 4-methoxyindol-3-ylmethyl. (B) LOD plot of relative methoxylated glucosinolate abundance in an Elbtalaue x Konstanz F2 population (C) Peak of the chromosome 2 quantitative trait locus (QTL), sorted by genotype. Mean +/− s.e. of N = 13 (Elbtalaue), 24 (Konstanz), and 45 (Heterozygote). (D) Peak of the chromosome 3 quantitative trait locus (QTL), sorted by genotype. Mean +/− s.e. of N = 36 (Elbtalaue), 10 (Konstanz), and 35 (Heterozygote). (E) Additive effects of the indole glucosinolate O-methylation based on the chromosome 2 and 3 genotypes. (F) LOD plot of fraction of modified indole glucosinolates in an Elbtalaue x Konstanz F2 population (G) Fraction of modified indole glucosinolates at 7 QTL, sorted by genotype. Mean +/− s.e. of N = 13 (Elbtalaue), 30 (Konstanz), and 40 (Heterozygote). Different letters indicates significant differences, P < 0.05, ANOVA followed by Tukey’s HSD test. Horizontal lines in panels B and F are 95% confidence levels, calculated based on 500 permutations of the data.

Arabidopsis has five indole glucosinolate methyltransferase (IGMT) genes. *IGMT1-4* (AT1G21100, AT1G21110, AT1G21120, and AT1G21130) are in a tandem-duplicated gene cluster on chromosome 1, and the more distantly related *IGMT5* (AT5G53810) is located on chromosome 5 [45,46]. Three predicted *E. cheiranthoides* IGMT genes (Erche01g022140.a, Erche01g022140.b, and Erche01g022140.d) are in a tandem-duplicated cluster on chromosome 1, and the encoded proteins are highly similar to the Arabidopsis IGMT1-4 (Figure 8A, Figure S9), which catalyze the O-methylation of 4HI3M to make 4MI3M. The most similar methyltransferases from *Raphanus sativus* (radish) and *B. oleracea* are shown for comparison in the phylogenetic tree. Consistent with the greater abundance of 4MI3M in Konstanz, two of the three *E. cheiranthoides* IGMT genes are expressed at a significantly higher level in Konstanz than in Elbtalaue (Figure 8B). In the F2 population, expression of all three *E. cheiranthoides* IGMT genes was positively correlated with the relative abundance of 4MI3M (Figure 8C,D,E). Quantitative trait mapping identified gene expression QTL on chromosome 6 for Erche01g022140.a, and on chromosome 3 for Erche01g022140.b and Erche01g022140.d (Figure S10). Chromosome 3 also has a QTL regulating the relative abundance of 4MI3M (Figure 7B), suggesting that *IGMT* gene expression variation may be the cause of the observed metabolite abundance QTL.

**Figure 8.**
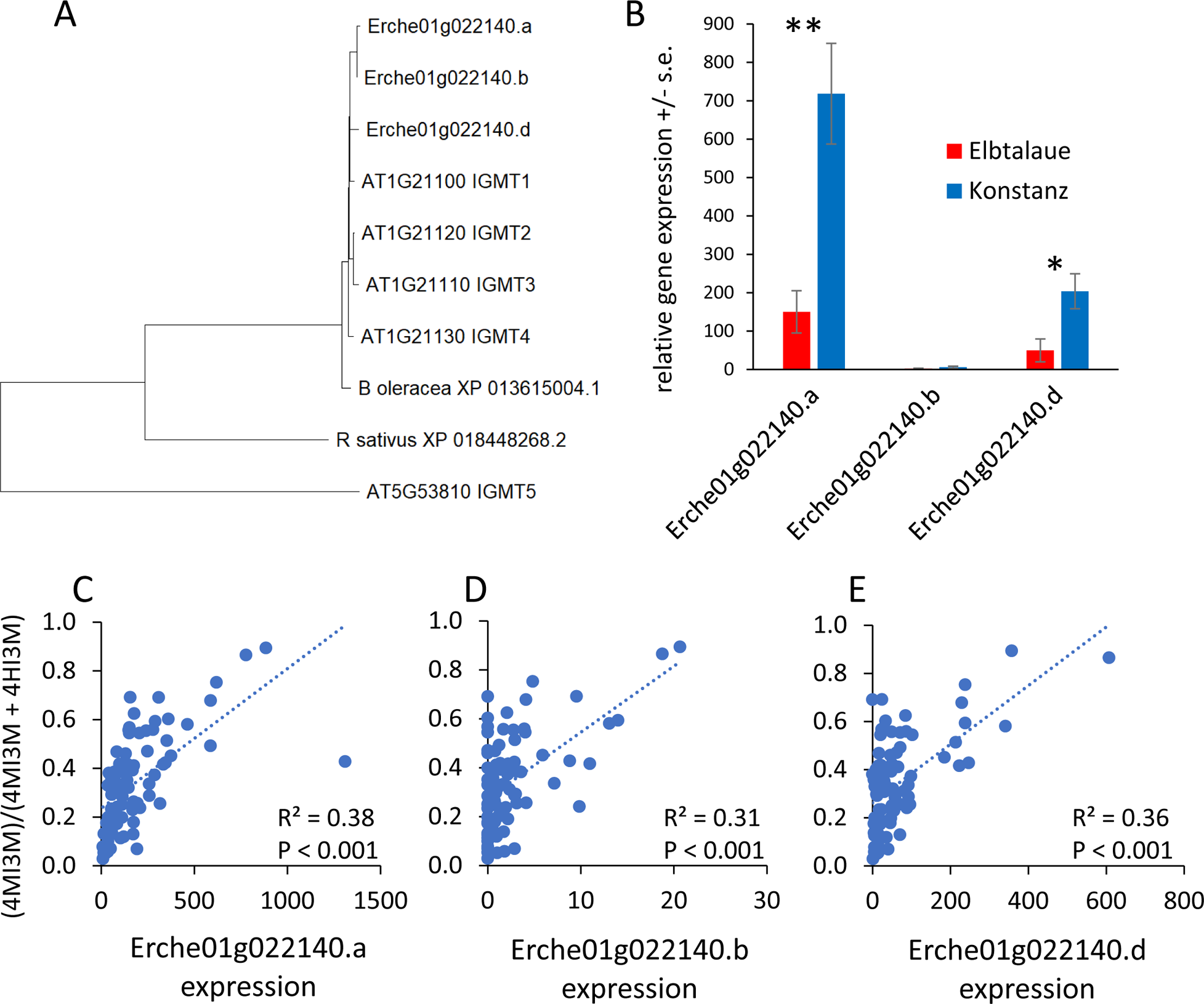
Expression of *Erysimum cheiranthoides* indole glucosinolate methyltransferase (IGMT) genes. (A) Neighbor-joining tree of predicted IGMT proteins from *E. cheiranthoides* (Erche), *Arabidopsis thaliana* (AT), radish (*Raphanus sativus*), and cabbage (*Brassica oleracea*), constructed using MEGA11. The corresponding protein sequence alignment is in Supplemental Figure S9. (B) Relative expression of *E. cheiranthoides IGMT* genes in the Elbtalaue and Konstanz lineages. * P < 0.05, **P < 0.005, *t-*test, N = 5, mean +/− s.e. (C, D, E) Correlation of *IGMT* expression (Erche01g022140.a, Erche01g022140.b, and Erche01g022140.d) and the abundance of methylated indole glucosinolates, (4MI3M)/(4MI3M + 4HI3M). 4HI3M = 4-hydroxyindol-3-ylmethylglucosinolate, 4MI3M = 4-methoxyindol-3-ylmethylglucosinolate. Lines show linear regression and P values are from Pearson correlations.

## 3. Discussion

By crossing two *E. cheiranthoides* inbred lines, we generated a segregating F2 population and used this to make a genetic map with 501 molecular markers (Figure S1). For the original *E. cheiranthoides* genome assembly, sequencing contigs were ordered into scaffolds using a Hi-C proximity ligation method [24]. Although this approach is efficient at placing assembled contigs in the right order on each chromosome, it is less reliable at placing contigs in the correct orientation. Based on the new genetic linkage map, we changed the relative orientations of individual contigs for several of the *E. cheiranthoides* chromosomes (Figure S3) and increased the percentage of the overall genome assembly that was anchored to chromosomes (Figure S4). This improved genome assembly not only made it possible to conduct reliable quantitative trait mapping for the current project but also will facilitate future genetic studies with *E. cheiranthoides*.

With the notable exception of helveticoside, there was significant positive correlation in the abundance of glucosinolates and cardiac glycosides in the F2 plants. Thus, there appears to be no major regulatory tradeoff in the production of these two classes of defensive metabolites in *E. cheiranthoides.* Among the detected cardiac glycosides and glucosinolates in our assays, only helveticoside was negatively correlated with aphid reproduction on plants in the F2 population (Figure 4A). When added to artificial diet, the IC50 of purified helveticoside was 14 ng/µl, which is similar to the 20 ng/mg wet weight concentration of this cardiac glycoside in *E. cheiranthoides* leaves [36]. However, it is not known at what concentration helveticoside is found in the phloem from which the aphids are feeding. We were not able to detect helveticoside in aphid tissue (Figure 3N), suggesting that it is either not localized in the phloem or somehow metabolized after it enters the aphids. However, the presence of helveticoside in aphids feeding on artificial containing this cardiac glycoside [36] suggests that complete conversion of helveticoside in aphids is less likely. Further research, ideally with mutations that specifically affect the production of helveticoside, will be needed to investigate the function of this metabolite in plant defense. A QTL affecting the abundance of helveticoside but not other cardiac glycosides (Figure 5) may lead to the eventual identification of biosynthetic or regulatory genes that specifically affect the production of this cardiac glycoside.

Homologs of known genes from Arabidopsis can account for most of the aliphatic glucosinolate biosynthesis pathway in *E. cheiranthoides* [23]. However, biosynthetic enzymes for glucosinolates that are present in *E. cheiranthoides* but not in Arabidopsis remain to be discovered. Leucine and isoleucine have both been described as amino acid precursors for glucosinolate biosynthesis [26] and could account for the structurally uncharacterized NMB glucosinolate, which is significantly more abundant in the Elbtalaue accession (Figure 2D). Cytochrome P450 enzymes in the CYP79F family have been associated with differential incorporation of methionine or branched-chain amino acids into *B. stricta* glucosinolates [41], and Erche01g017900, a gene encoding a predicted CYP79F enzyme, is within the confidence interval of an NMB QTL on chromosome 1 (Figure 6A).

The expression levels of the Elbtalaue and Konstanz alleles of Erche01g017900 were not significantly different (Figure S6B), and there is only one amino acid sequence difference between the two accessions, glycine and serine, respectively, at position 51 (Figure S6A). *Bs*BCMA1 and *Bs*BCMA3, two *B. stricta* enzymes that preferentially catalyze the incorporation of branched-chain amino acids rather than methionine into glucosinolates [41], have a serine this position, whereas *Bs*BCMA2 has a glycine (Figure S6). Since Konstanz has the serine allele at position 51, differences in Erche01g017900 enzymatic activity may not explain the lower NMB abundance relative to Elbtalaue (Figure 2D).

Biosynthetic enzymes for methylsulfonyl glucosinolates have not yet been identified in any plant species. A family of flavin-dependent monooxygenases catalyze the formation of melthylsulfinyl glucosinolates in Arabidopsis [47,48], and it is possible that similar enzymes catalyze further oxidation of glucosinolate substrates to produce methylsulfonyl glucosinolates in *E. cheiranthoides.* Both genetic mapping (Figure 6C) and analysis of genes with expression patterns that are similar to those encoding other aliphatic glucosinolate biosynthetic enzymes (Figure 6E) may lead to the identification of such enzymes in *E. cheiranthoides*.

In Arabidopsis, three CYP81F monooxygenases (AT4G37430, AT4G37400, and AT5G57220) and four IGMTs (AT1G21100, AT1G21110, AT1G21120, and AT1G21130) [44,45] catalyze the sequential modification of I3M to form 4HI3M and 4MI3M (Figure 7A). Formation of hydroxylated and methoxylated indole glucosinolates is induced as a defense response, and the presence of multiple enzymes with similar functions may allow more complex regulation of this process. To accomplish this, the multiple indole glucosinolate modifying enzymes may be subject to differential regulation. Among the two IGMTs that are expressed at a significantly higher level in Konstanz (Figure 8B), expression of Erche01g022140.a is regulated by a QTL on chromosome 6 and expression of Erche01g022140.c is regulated by a QTL on chromosome 3.

A QTL on chromosome 7 (Figure 7F) may be associated with increased I3M hydroxylase activity. However, the *E. cheiranthoides* homologs of Arabidopsis CYP81F monooxygenases that catalyze I3M 4-hydroxylation [44,45], are encoded on chromosome 2 (Erche02g041710 and Erche02g041680). Therefore, cis-regulation or differences in enzymatic activity are unlikely to be the cause of this variation in the indole glucosinolate profile. Moreover, Erche02g041710 and Erche02g041680 do not have significant expression QTL on chromosome 7, the location of a QTL affecting the (4MI3M + 4HI3M)/(I3M + 4MI3M + 4HI3M) ratio, indicating that this QTL does not effect the expression of Erche02g041710 and Erche02g041680.

Despite the the significanty higher aphid reproduction on Elbtalaue than on Konstanz (Figure 1B), genetic mapping of this trait in an F2 population identified no significant QTL. A likely explanation is that there are multiple loci affecting aphid resistance, none of which have an effect that is large enough to be identified in an F2 population with only 83 genotyped plants. The hypothesis of multiple loci independently causing aphid resistance is also consistent with the observation that aphid survival and reproduction are not significantly different between Elbtalaue and F2 plants (Figure 1A,B). For instance, if multiple R-genes from the Elbtalaue genotype independently cause dominant resistance in the F2 population, aphid performance on the average F2 plant would be similar to Elbtalaue. R-gene mediated resistance to aphids has been observed in other plant species, including tomato and melon [49,50].

Differences in cardiac glycoside abundance do not adequately explain the improved performance of aphids on Elbtalaue relative to Konstanz plants. Although cheirotoxin, erysimoside, erychroside, and glucodigitoxigenin were more abundant in the Elbtalaue accession (Figure 3A-D), abundance of these cardiac glycosides was not negatively correlated with aphid performance on F2 plants (Figure 4A). Conversely, although helveticoside abundance is negatively correlated with aphid resistance in the F2 population (Figure 4A), there is no significant difference in the abundance of this compound between the two parent lines (Figure 3K). The greater helveticoside variation in the F2 population is due to transgressive segregation, and the lack of helveticoside is unlikely to be the cause of improved aphid growth on Konstanz plants. Performance of *M. persicae* also was not significantly improved on *cyp87a126* mutant *E. cheiranthoides* plants, which have a complete knockout of cardiac glycoside biosynthesis [15].

Although aphid feeding did not induce overall glucosinolate accumulation, aphids feeding on Konstanz plants for 24 h had elevated levels of 4MI3M in their bodies (Figure 2F) relative to earlier timepoints, suggesting increased abundance of this compound in the phloem. In Arabidopsis experiments, indole glucosinolate breakdown products were aphid-deterrent [37], and induced 4MI3M accumulation increased aphid resistance [11]. However, 4MI3M abundance was positively correlated with aphid reproduction in the *E. cheiranthoides* F2 population (Figure 4A). Given that 4MI3M abundance is positively correlated with other *E. cheiranthoides* metabolites, it is possible that additional defenses mask the predicted negative effects of 4MI3M. It is also possible that other factors in *E. cheiranthoides* influence the breakdown of 4MI3M and make this compound less toxic in this experimental context than when aphids consume 4MI3M from Arabidopsis.

Together, experiments with our *E. cheiranthoides* F2 population have resulted in an improved genome assembly and new insights into the biosynthesis and defenseive functions of glucosinolates and cardiac glycosides. Although aphid reproductive fitness, cardiac glycoside content, and glucosinolate content all vary between the two parental lines of the F2 population, variation in the abundance of the two classes of defensive metabolites do not adequately explain the observed differences in aphid performance. This indicates that additional, as yet unknown mechanisms of aphid resistance exist in *E. cheiranthoides.* A diverse defensive repertoire likely provides benefits in defense against generalist herbivores like *M. persicae* that are relatively tolerant of both glucosinolates and cardiac glycosides.

## 4. Materials and Methods

### 4.1 Plant and insect rearing

*Erysimum cheiranthoides* accession Elbtalaue, which has a published genome sequence [24], was collected in the Elbe River floodplain (Elbtalaue) in Lenzen, Germany. The Konstanz accession was originally collected in Oggenhausen, Germany, and was propagated at the Konstanz Botanical Garden in Konstanz, Germany. Seed stocks of both *E. cheiranthoides* accessions are available from the Arabidopsis Biological Resource Center (www.arabidopsis.org; stock numbers CS29250 and CS29251, respectively). We grew all plants in Cornell Mix [by weight 56% peat moss, 35% vermiculite, 4% lime, 4% Osmocote slow-release fertilizer (Scotts, Marysville, OH), and 1% Unimix (Scotts, Marysville, OH)] in 6 × 6 × 6-cm pots in a Conviron (Winnipeg, Canada) growth chamber, with 200 mmol m^−2^ s^−1^ light intensity at 23°C, with 50% relative humidity and a 16h/8h day/night cycle. We conducted all insect assays with a tobacco-adapted *M. persicae* strain [51–53] that we maintained on *Nicotiana tabacum* (tobacco) with 150 mmol m^−2^ s^−1^ light intensity at 24/19 °C day/night temperature, with 50% relative humidity and a 16h/8h day/night cycle.

### 4.2 Insect bioassays

For aphid survival and reproduction assays, we placed groups of five fourth-instar *M. persicae* into clip cages on *E. cheiranthoides* leaves. After 10 days, we counted the number of surviving adult aphids and nymphs. For the time-series aphid experiment, we utilized five-week-old *E. cheiranthoides* plants, with each plant hosting a group of 15 fourth-instar *M. persicae* aphids enclosed within clip cages. Aphids along with the leaf areas surrounded by these cages were collected in separate tubes and promptly frozen in liquid nitrogen after 1, 8, and 24 hours. For aphid choice assays, we placed one leaf each of Elbtalaue and Konstanz plants into 15 cm diameter Petri dishes, with their petioles inserted into a piece of moistened filter paper. To determine aphid feeding preferences, we released 10 adult *M. persicae* at the midpoint between the two leaves and, 24 hours later, counted the number of aphids on each leaf. Aphids that were not on either of the two leaves were not included in the data analysis. For artificial diet assays, we assembled aphid cages with 200 µl artificial diet [54,55], containing helveticoside (www.cfmot.de) at concentrations ranging from 0 to 100 ng/µl, between two layers of stretched Parafilm at the top of the cage. We placed 10 adult aphids into each cage and, after 7 days, we counted the number of surviving aphids and progeny in each cage. The experiment was conducted with 4 replicates.

### 4.3 Detection of glucosinolates and cardiac glycosides

For measurement of glucosinolates and cardiac glycosides, we prepared methanol extracts of *E. cheiranthoides* leaves and whole aphids, and analyzed them by HPLC-MS, as described previously [36]. Eight glucosinolates [indol-3-ylmethylglucosinolate (I3M) 4-hydroxyindol-3-ylmethylglucosinolate (4HI3M), 4-methoxyindol-3-ylmethylglucosinolate (4MI3M), 3-methylsulfinylpropylglucosinolate (3MSIP), 3-methylsulfonylpropylglucosinolate (3MSOP), 4-methylsulfonylbutylglucosinolate (4MSOB), 3-methylthiopropylglucosinolate (3-MTP), and n-methylbutylglucosinolate (NMB, a structurally uncharacterized glucosinolate with a 5-carbon side chain)] and 7 cardiac glycosides [cheirotoxin, erysimoside, erychroside, glucodigitoxigenin, helveticoside, erycordin, and dig-10 (a digitoxigenin-derived cardiac glycoside with a structurally uncharacterized sugar)] were detected in these assays.

### 4.4 Transcriptome sequencing

We sequenced the transcriptomes of the 83 F2 individuals from a cross between *E. cheiranthoides* accessions Elbtalaue and Konstanz using the 3’RNAseq method [56]. Additionally, we sequenced RNA from 5 Elbtalaue and 5 Konstanz samples, which served as parental references. RNA was isolated from frozen harvested materials using the SV Total RNA isolation kit with on-column DNA digestion (Promega, Madison, WI, USA). The purity of all RNA samples was confirmed using a NanoDrop2000 instrument (Thermo Scientific). The 3’RNA-seq libraries were prepared from 6 µg total RNA at the Cornell Genomics facility (http://www.biotech.cornell.edu/brc/genomics-facility) [56]. Transcriptome sequencing data were deposited in the Sequence Read Archive (https://www.ncbi.nlm.nih.gov/sra) under accession PRJNA1053801.

### 4.5 Genetic map construction and assembly of E. cheiranthoides genome v2.0

We performed read mapping and SNP calling by following the Genome Analysis ToolKit (GATK) best practices for RNAseq short variant discovery [57,58]. 3’RNAseq data from 83 F2 plants, five var. Konstanz, and five var. Elbtalaue plants were aligned to unpolished PacBio contigs using STAR version 2.7.1a default parameters and 2-pass mapping [59]. The resulting bam files were cleaned using GATK tools MarkDuplicates, AddOrReplaceReadGroups, and SpljitNCigarReads. Variants were called with HaplotypeCaller, and joint genotyping was performed using GenotypeGVCFs [60]. The resulting VCF file was filtered using bcftools filter [61] to include only biallelic SNPs with a quality score greater than 30, alternate allele frequency between 0.3-0.7, excess heterozygosity less than two, and a called genotype in at least half of the samples. The filtered VCF was converted to ABH using Tassel 5 [62], the markers were binned using SNPbinner [63], and a genetic map was made using MSTmap [64]. During map construction, one contig was found to be chimeric and was split at the most likely splice point, as determined by a visual analysis of aligned PacBio reads. The resulting genetic map was reconciled with the Hi-C proximity guided assembly [23,24] using a custom Python script (https://github.com/gordonyounkin/Erysimum_F2_aphids) that prioritized placement and orientation of contigs in the genetic map. The final fasta assembly containing pseudomolecules and contigs was constructed using CombineFasta (https://github.com/njdbickhart/CombineFasta). Illumina reads were aligned to the new genome using Burrows-Wheeler Aligner version 0.7.8 [65], and the assembly was polished with three rounds of Pilon version 1.23 [66]. The chloroplast genome was assembled from PacBio reads using Organelle_PBA [67,68]. Plots were generated in R [69] using R/qtl [70].

### 4.6 Genome annotation

Gene annotations were transferred from version 1.2 to version 2.0 of the *E. cheiranthoides* genome using GMAP [71]. Annotations were improved by aligning full length *E. cheiranthoides* RNA sequencing reads (NCBI: PRJNA563696) to the new genome assembly with hisat2 [72], sorting aligned reads with samtools [73], and assembling and merging transcripts with StringTie [74]. In cases where there was not a 1:1 relationship between stringtie transcripts and the original gene annotations, a new name was assigned to each transcript. Open reading frames and protein sequences were predicted using getorf from EMBOSS [75].

### 4.7 Coexpression network analysis

RNA sequencing reads from the *E. cheiranthoides* F2 population were pseudoaligned to the transcriptome associated with *E. cheiranthoides* genome version 2.0 (NCBI: PRJNA563696) [76] using kallisto with default parameters, yielding transcript counts [77]. Transcripts with more than 10 counts in at least 50 samples were retained for either dataset. Filtered counts were used for the mr2mods gene coexpression analysis pipeline using default parameters [42] (https://github.itap.purdue.edu/jwisecav/mr2mods). Predicted *E. cheiranthoides* glucosinolate biosynthesis genes [23], were used as baits to identify network modules related to glucosinolate biosynthesis. Co-expression networks were visualized using using Cytoscape v3.9.1 (https://cytoscape.org) [78].

### 4.8 Data analysis

ANOVA and *t-*tests were conducted using JMP Pro 16 (https://www.jmp.com). We calculated the IC50 (cardiac glycoside concentration to reduce progeny production by 50%) using Solver function in Excel to fit a curve of the form: Y = 1/(1 + exp [B − G · ln(X)]), where X is cardiac glycoside concentration, Y is the fraction of larvae killed by the infection, and B and G are parameters which are varied for optimal fit of the curve to the data points (minimizing the residuals). We conducted QTL mapping using Windows QTL Cartographer [79]. Sequences were aligned using Clustal Omega [80]. Neighbor joining trees were constructed using default parameters in MEGA11 [81]. For Pearsson correlations of metabolite and aphid resistance data, the data were transformed to normality using a two-step process in SPSS (https://www.ibm.com), as described previously [82]. Raw data underlying all manuscript figures are included in Tables S2-S11.

## Supplementary Materials

The following supporting information can be downloaded at: www.mdpi.com/xxx/s1

**Figure S1.** Distribution of 501 genetic markers in the Konstanz x Elbtalaue genetic map across 8 chromosomes

**Figure S2.** Pairwise recombination fractions and LOD scores indicating probability of genetic linkage for 501 genetic markers

**Figure S3**. Position of genetic markers in *E. cheiranthoides* genome versions 1.2 and 2.0

**Figure S4.** Comparison of assembly statistics for *E. cheiranthoides* genome versions 1.2 and 2.0

**Figure S5.** Allele frequencies across the F2 mapping population

**Figure S6**. Alignment of CYP79F sequences

**Figure S7.** Neighbor joining tree of CYP79F protein sequences

**Figure S8.** QTL affecting oxygenated sulfur glucosinolate abundance

**Figure S9.** Alignment of indole glucosinolate methyltransferase proteins sequences

**Figure S10.** QTL affecting indole glucosinolate methyltransferase gene expression

**Table S1.** Coexpression network with aliphatic glucosinolate biosynthesis genes

**Tables S2-S11.** Raw data underlying manuscript figures

## Author Contributions

Conceptualization, GJ and MM; Methodology, MM, GCY, and GJ; Software, GCY and AFP; Validation, MM and GCY; Formal Analysis, MM, GCY, AFP, MLA, and GJ; Investigation, MM, GCY, AFP, and MLA; Resources, GJ; Data Curation, GCY; Writing – Original Draft Preparation, GJ; Writing – Review & Editing, MM, GCY, and GJ; Visualization, MM, GCY, and GJ; Supervision, SRS and GJ; Project Administration, GJ; Funding Acquisition, GJ, GCY, and MLA.

## Data Availability Statement

Version 2.0 of the *E. cheiranthoides* genome is available from GenBank (accession number PRJNA563696), and an annotated version of the genome is available at www.erysimum.org. Transcriptome sequencing data generated through this research have been deposited in GenBank (accession number PRJNA1053801). Raw data underlying figures in this manuscript are presented in the Supplemental Tables S2-S11.

---

**Figure S1.**
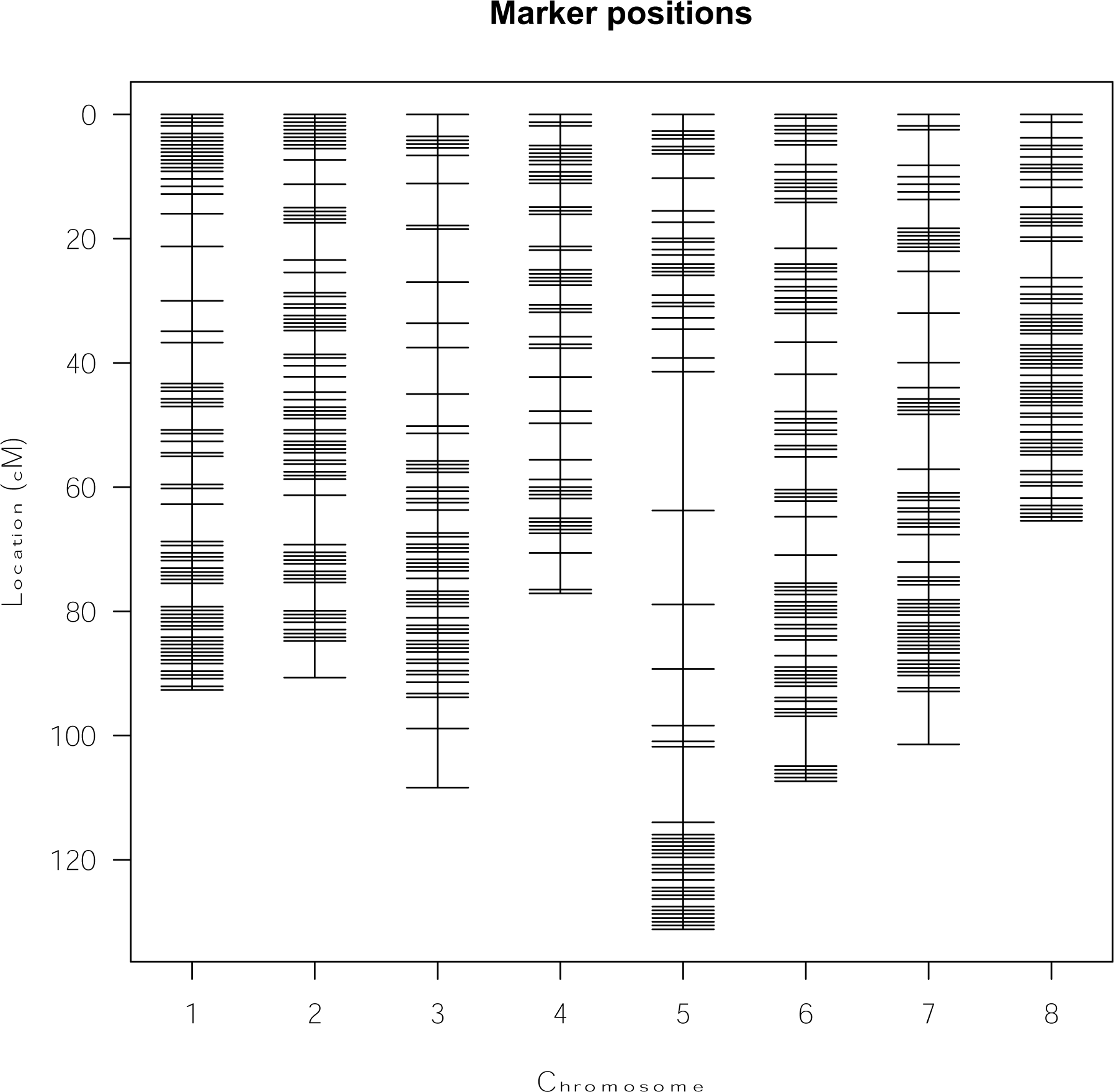
Distribution of 501 genetic markers in the *Erysimum cheiranthoides* Konstanz x Elbtalaue F2 population genetic map. Figure was generated using the plotMap() function in R/qtl.

**Figure S2.**
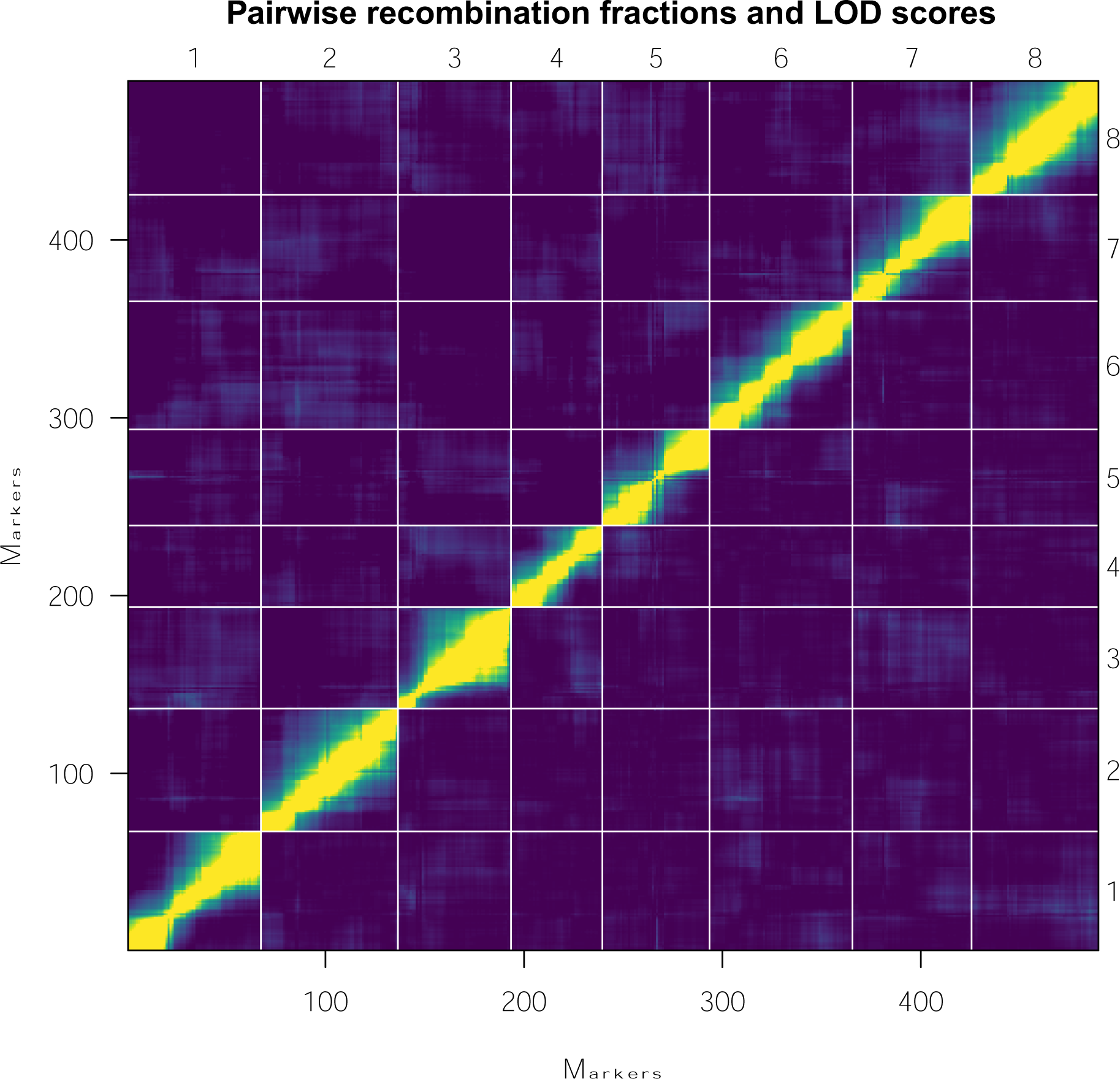
Genetic linkage of *Erysimum cheiranthoides* chromosomal markers. Pairwise recombination fractions (above diagonal) and LOD scores (below diagonal) indicating probability of genetic linkage for 501 genetic markers in the Konstanz x Elbtalaue genetic map. The 8 linkage groups correspond to the 8 chromosomes of *E. cheiranthoides*. Figure was generated using the plotRF() function in R/qtl.

**Figure S3.**
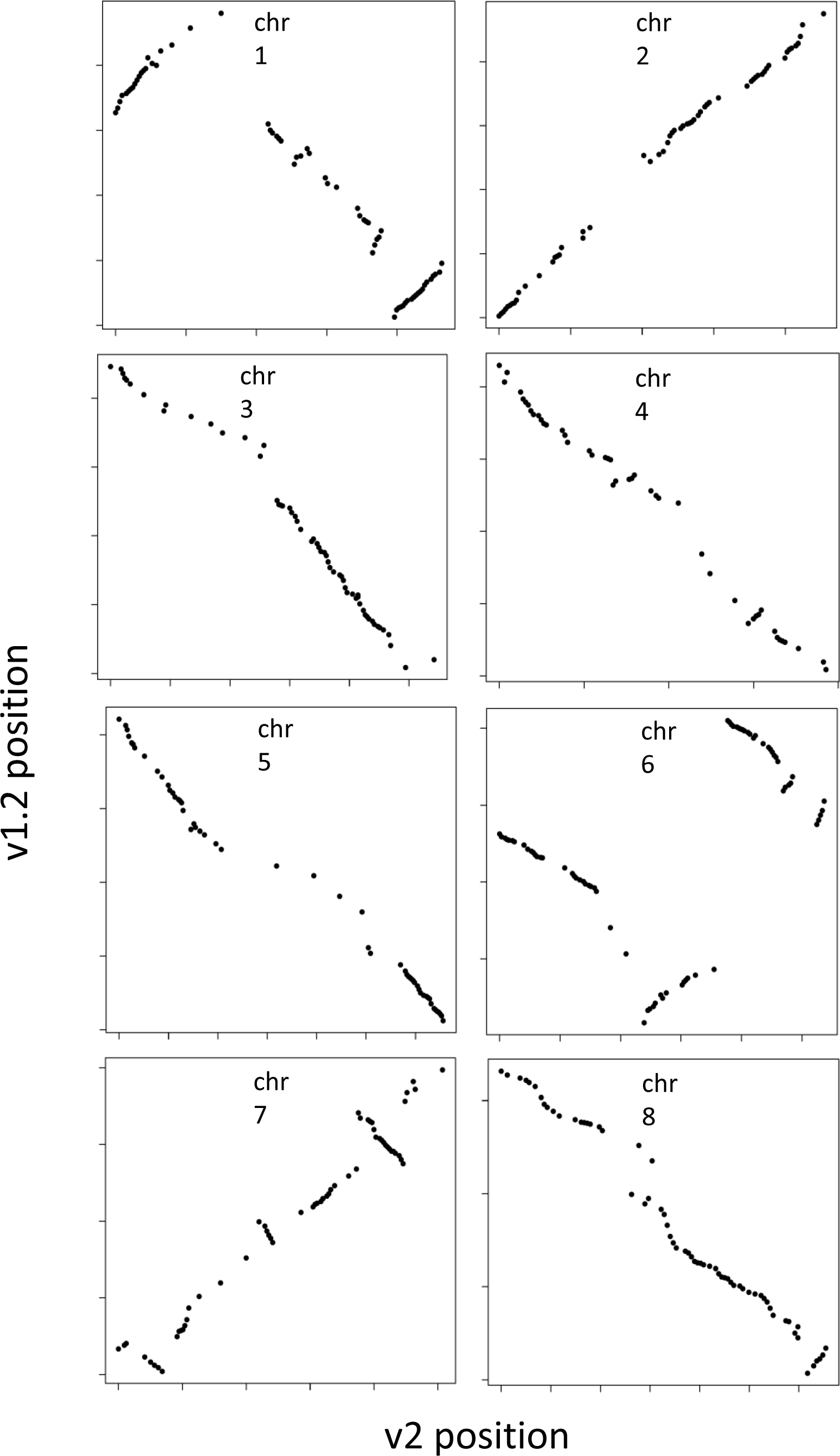
Position of genetic markers in *Erysimum cheiranthoides* genome v1.2 (Hi-C proximity guided assembly) and v2.0 (assembled using a classical genetic map). Large regions on chromosomes 1, 4, 6, 7, and 8 were inverted or incorrectly ordered in v1.2.

**Fiugre S4.**
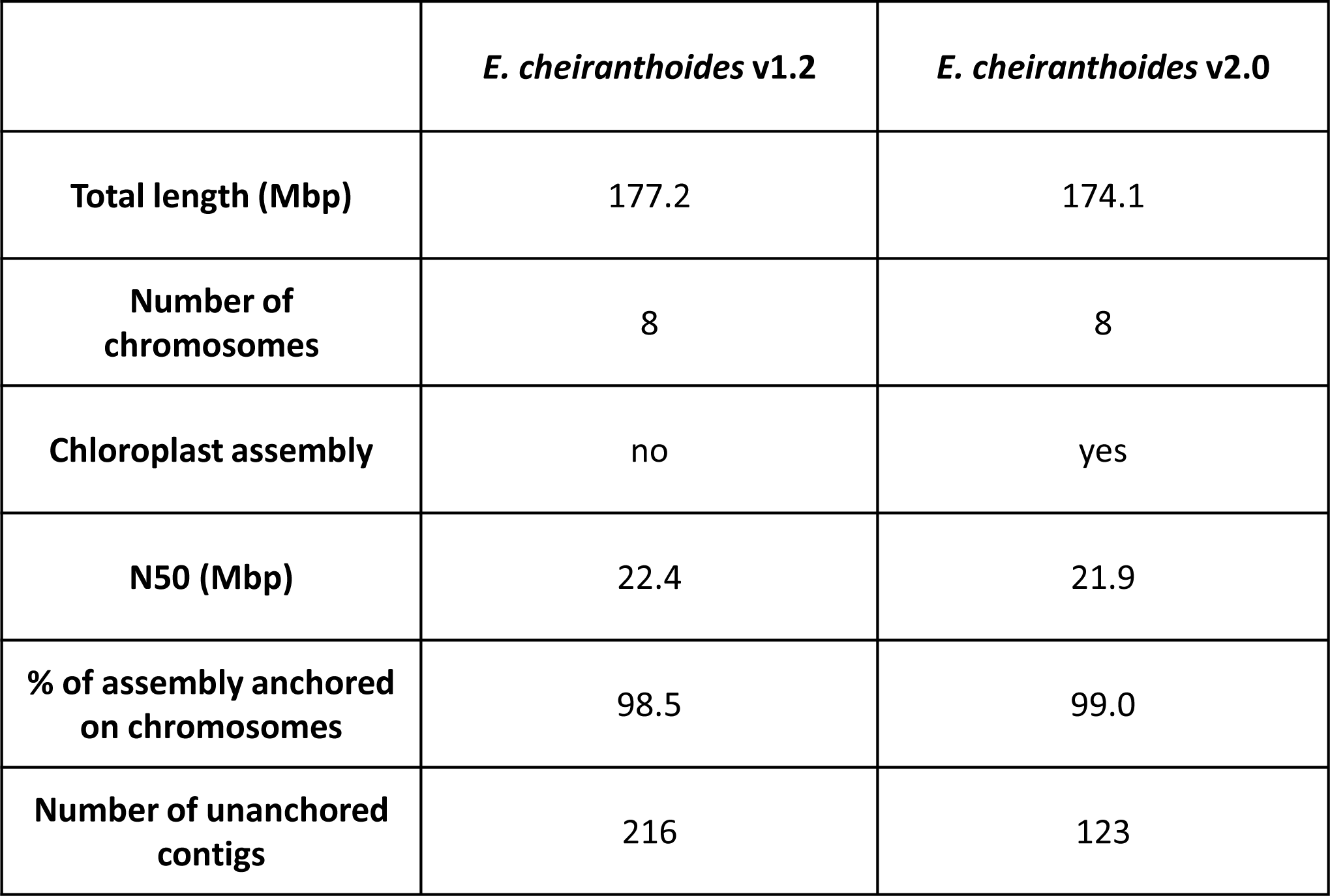
Comparison of assembly statistics for *Erysimum cheiranthoides* genome v1.2 and v2.0. In addition to improved congruency with genetic mapping data (Figure 3), v2.0 includes a chloroplast assembly and fewer unanchored contigs. It is unclear why v2.0 is 3.1 Mbp shorter than v1.2, but the total length of v2.0 matches the total length of the PacBio contigs.

**Figure S5.**
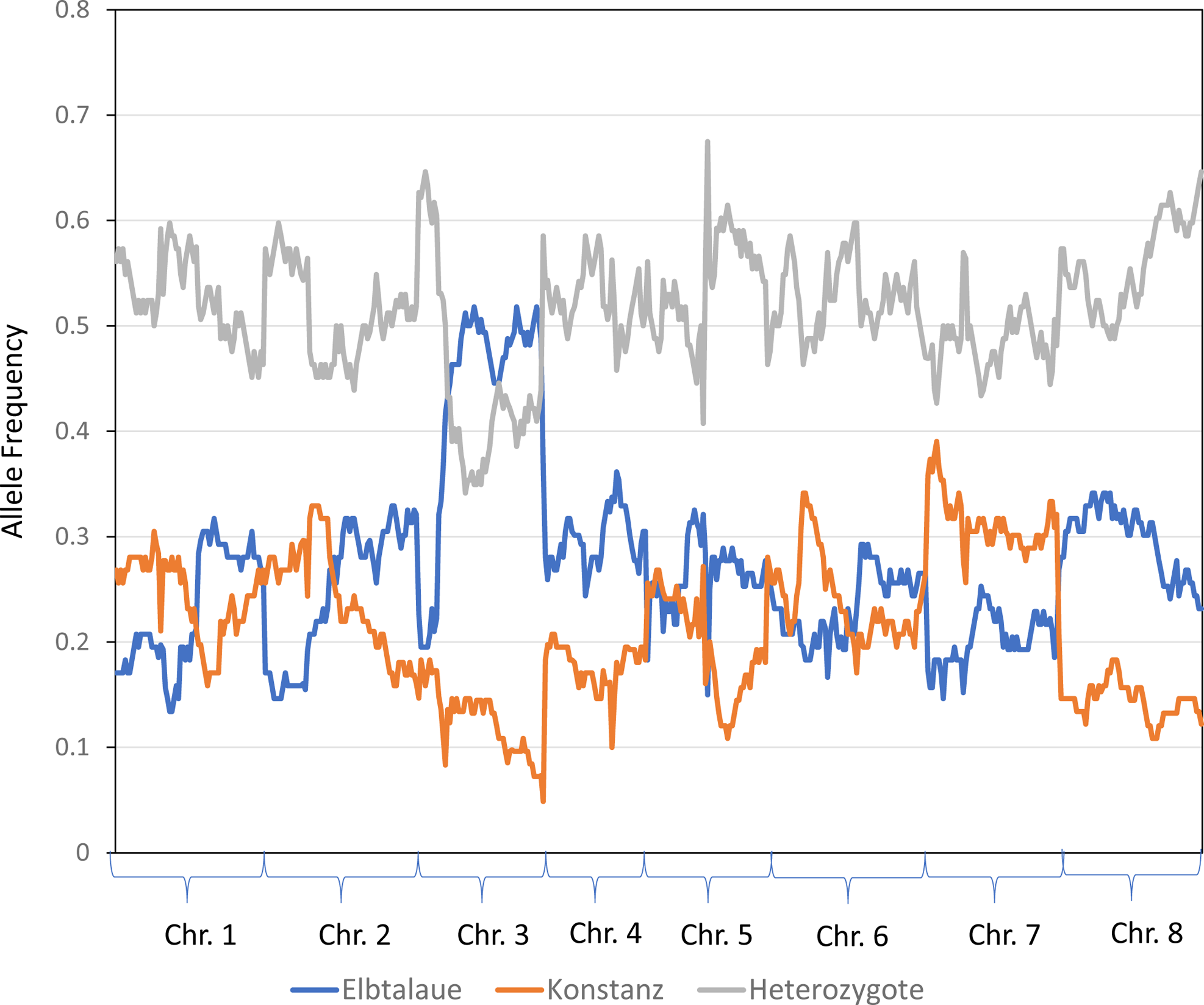
Allele frequencies across the *Erysimum cheiranthoides* F2 mapping population. The frequency of homozygous Elbtalaue, homozygous Konstanz, and heterozygous alleles for 501 molecular markers across 83 F2 lines is shown.

**Figure S6.**
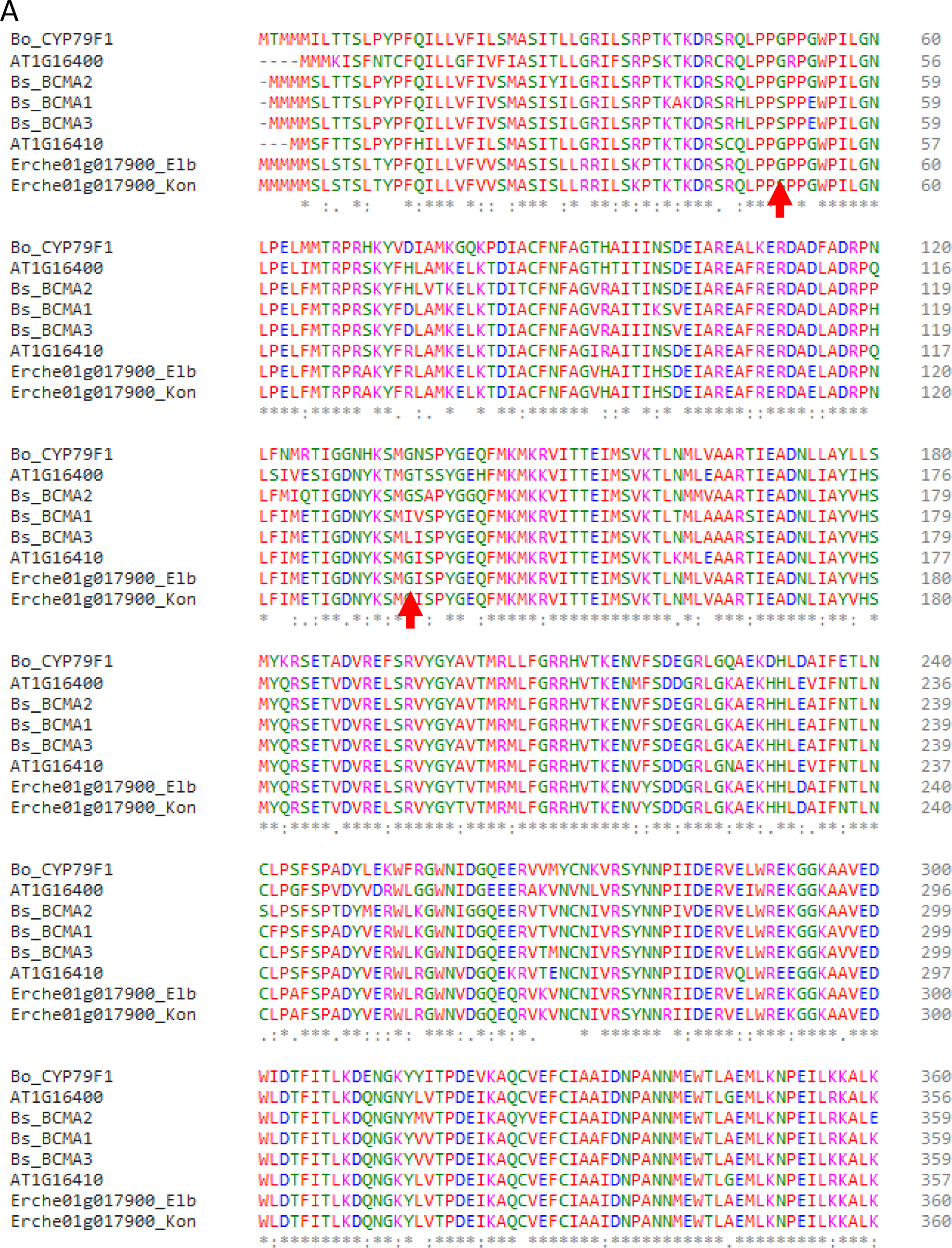
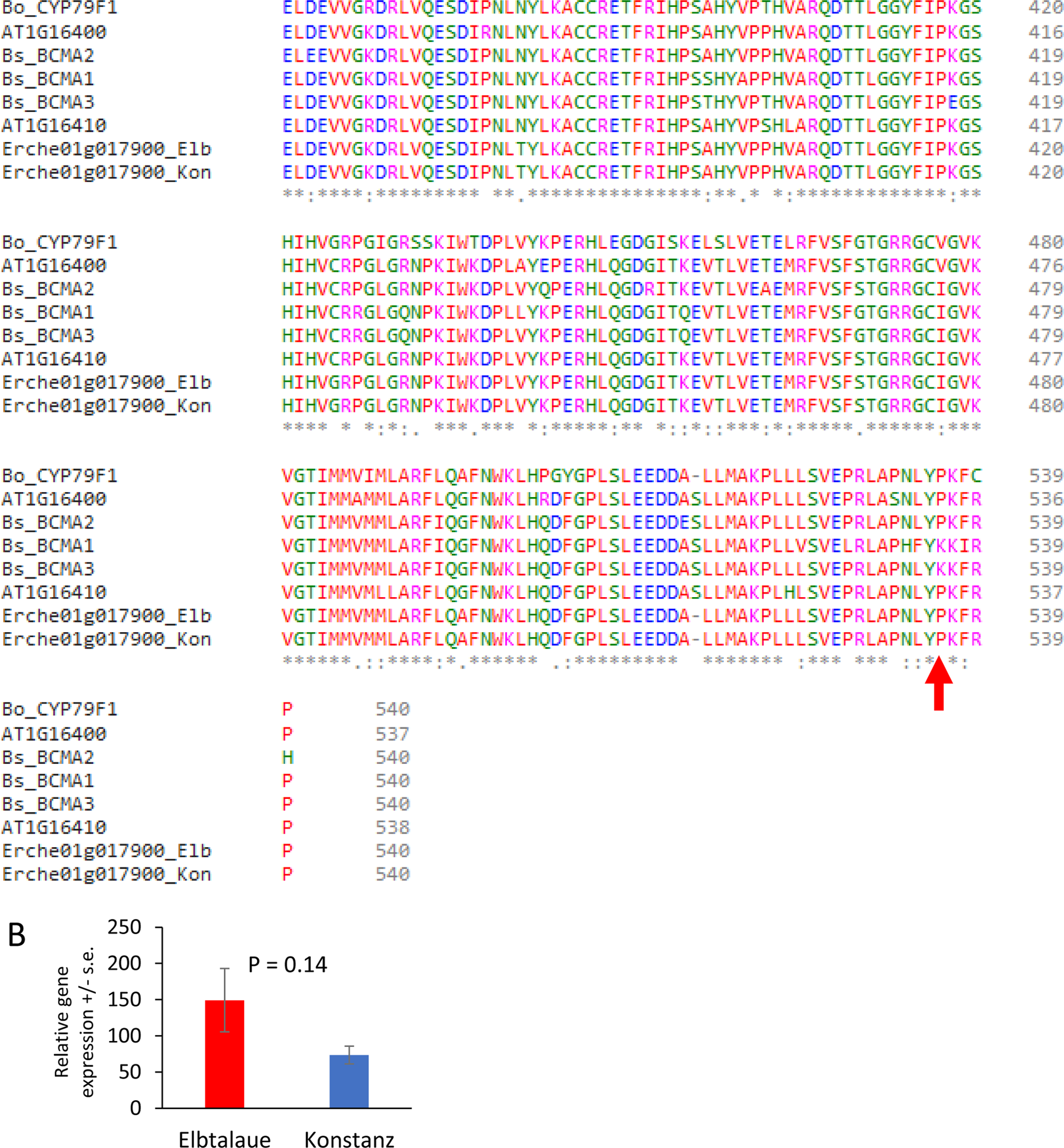
Alignment of CYP79F sequences and gene expression. (A) CYP79F protein sequences from *Erysimum cheiranthoides* (Erche), *Arabidopsis thaliana* (AT)*, Boechera stricta* (Bs), and *Brassica oleracea* (Bo) were aligned using Clustal Omega (https://www.ebi.ac.uk/Tools/msa/clustalo/). Red arrows indicate positions with amino acid variation in *B. stricta* that has been associated with the differences in the preferential incorporation of methionine (BCMA2) or branched chain amino acids (BCMA1 and BCMA3) into glucosinolates side chains (Prasad et al, 2012, *Science* 337:1081-1084). The *E. cheiranthoides* Elbtalaue (Elb) and Konstanz (Kon) alleles differ at one of these positions, having glycine and serine, respectively at position 51. (B) Expression levels of Erche01g017900 in the Elbtalaue and Konstanz accessions. Mean +/− s.e. of N = 5; P value is from a two-tailed *t-*test.

**Figure S7.**
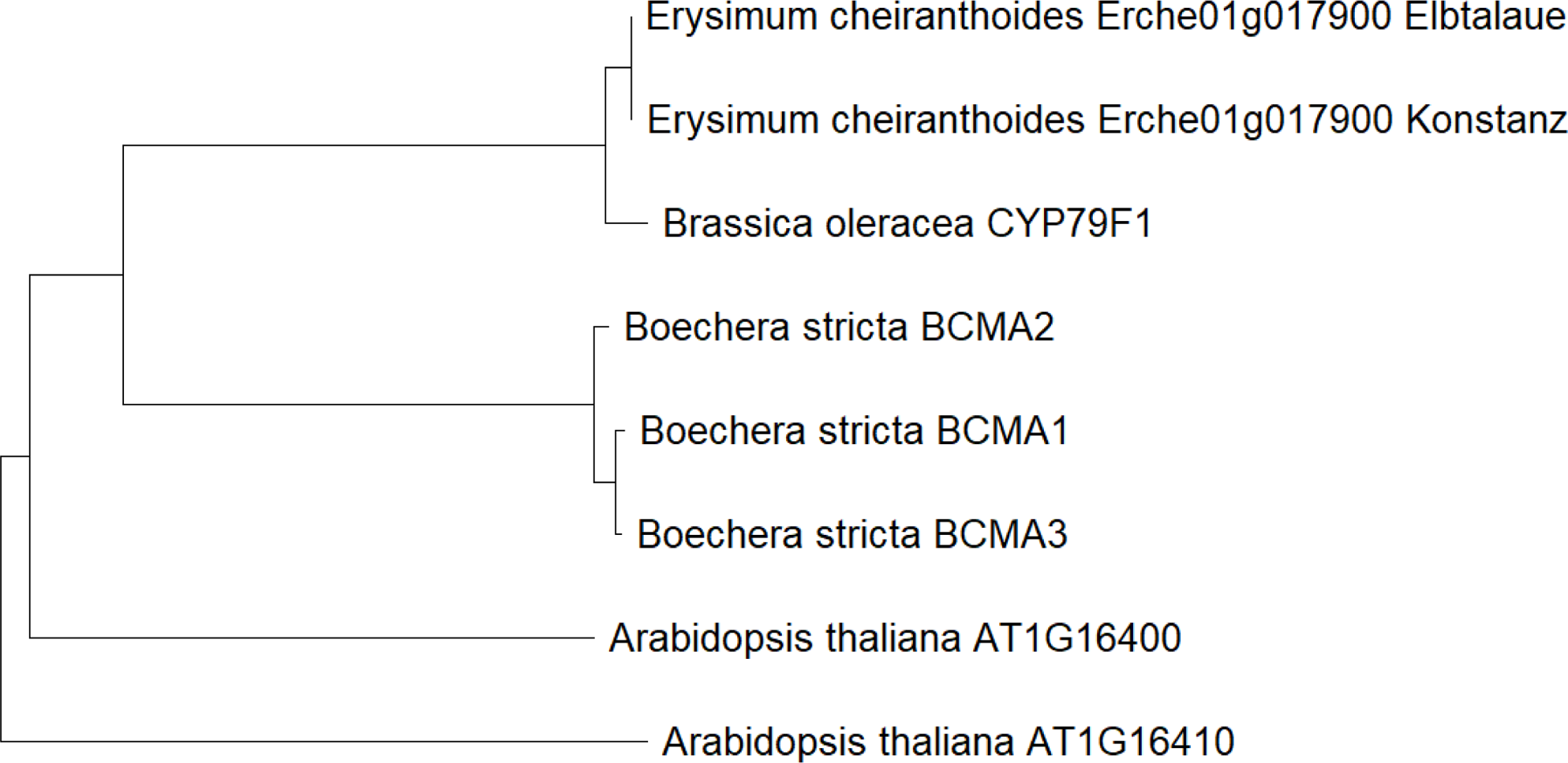
Neighbor-joining tree of CYP79F protein sequences. CYP79F protein sequences from *Erysimum cheiranthoides*, *Arabidopsis thaliana*, *Boechera stricta*, and *Brassica oleracea* were aligned using the Muscle algorithm and a neighbor-joining tree was constructed using MEGA11.

**Figure S8.**
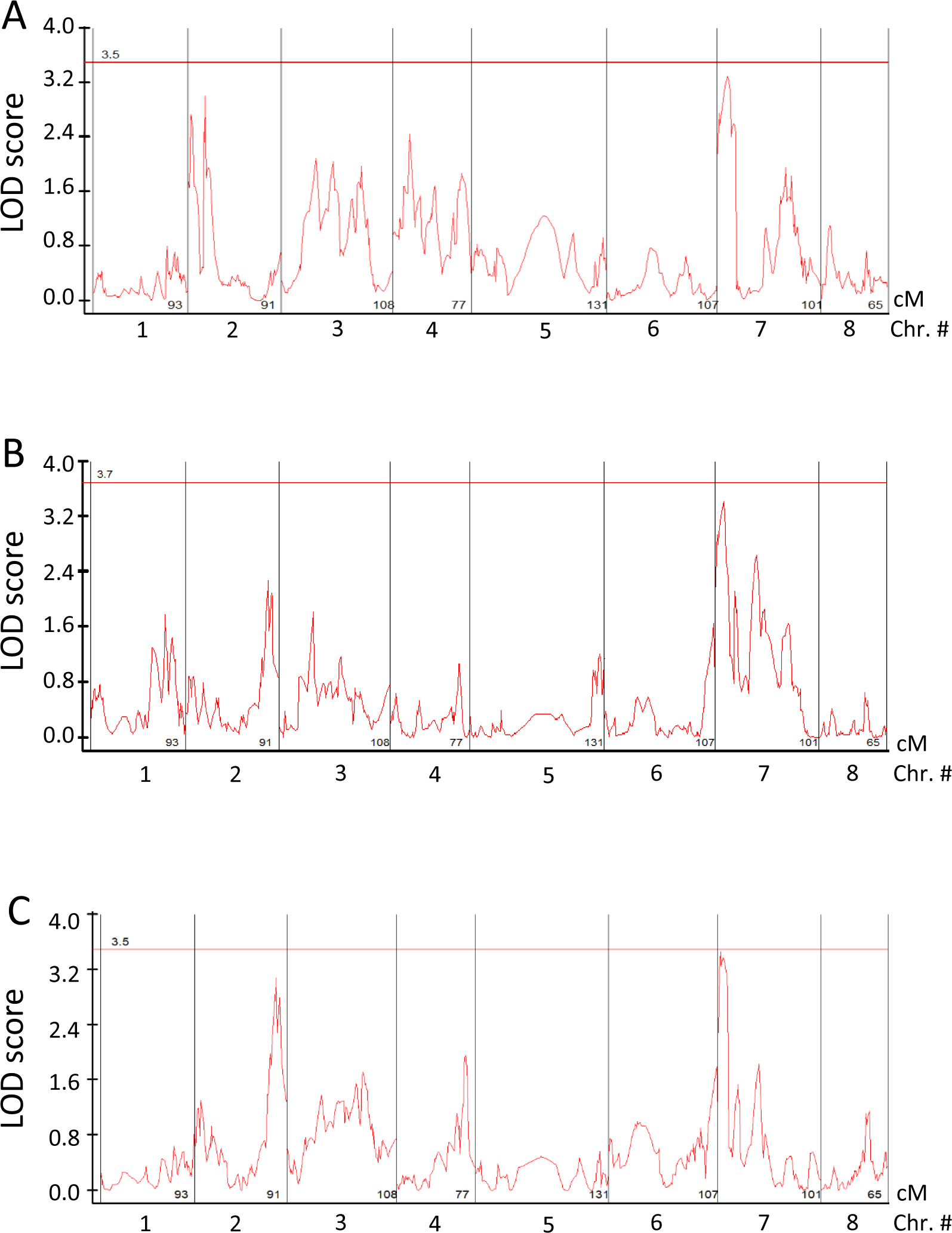
Quantitative trait loci (QTL) affecting glucosinolate abundance in *Erysimum cheiranthoides*. LOD plots of (A) 3-methylsulfinylpropylglucosinolate (3MSIP), (B) 3-methylsulfonylpropylglucosinolate (3MSOP), and (C) 4-methylsulfonylbutylglucosinolate (4MSOB) abundance in an Elbtalaue x Konstanz F2 population are shown. Horizontal lines in are 95% confidence levels, calculated based on 500 permutations of the data. Graphs were created using Windows QTL Cartographer.

**Figure S9.**
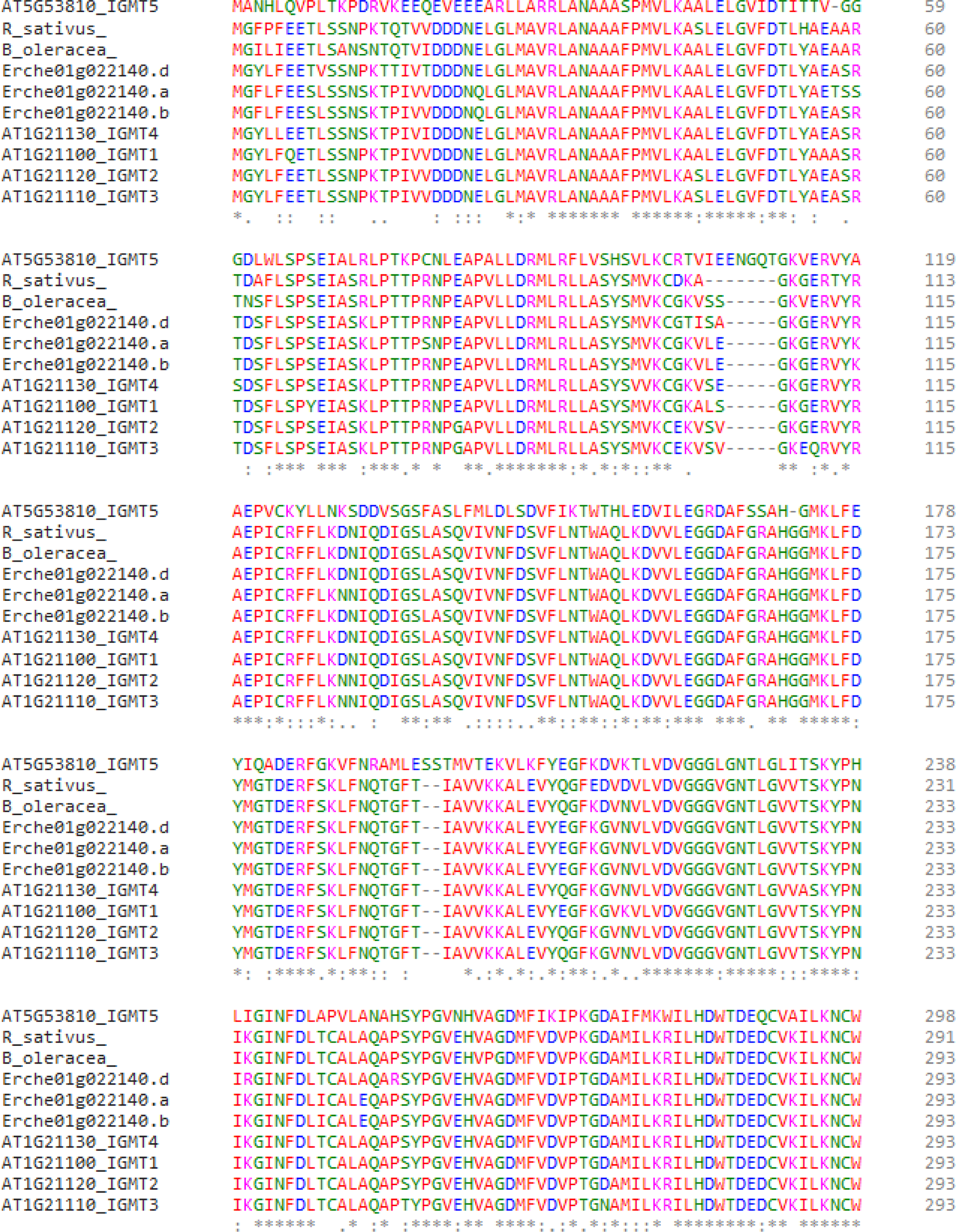
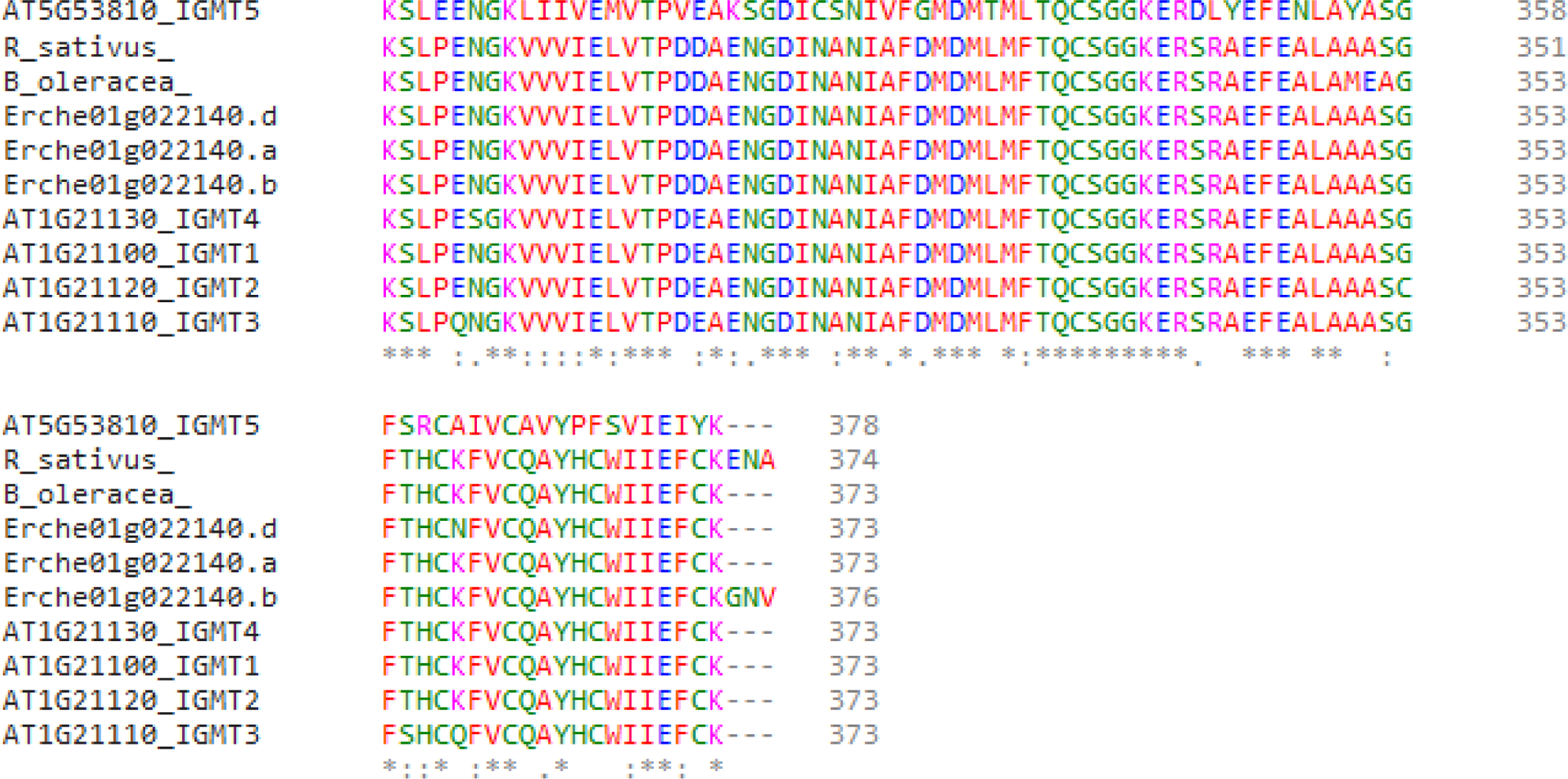
Alignment of indole glucosinolate methyltransferase (IGMT) protein sequences. Sequences of predicted IGMT proteins from *Erysimum cheiranthoides* (Erche01g022140.a, Erche01g022140.b, and Erche01g022140.d), *Arabidopsis thaliana* (AT1G21100, AT1G21110, AT1G21120, AT1G21130, and AT5G53810), radish (*Raphanus sativus*), and cabbage (*Brassica oleracea*) were aligned using Clustal Omega (https://www.ebi.ac.uk/Tools/msa/clustalo/)

**Figure S10.**
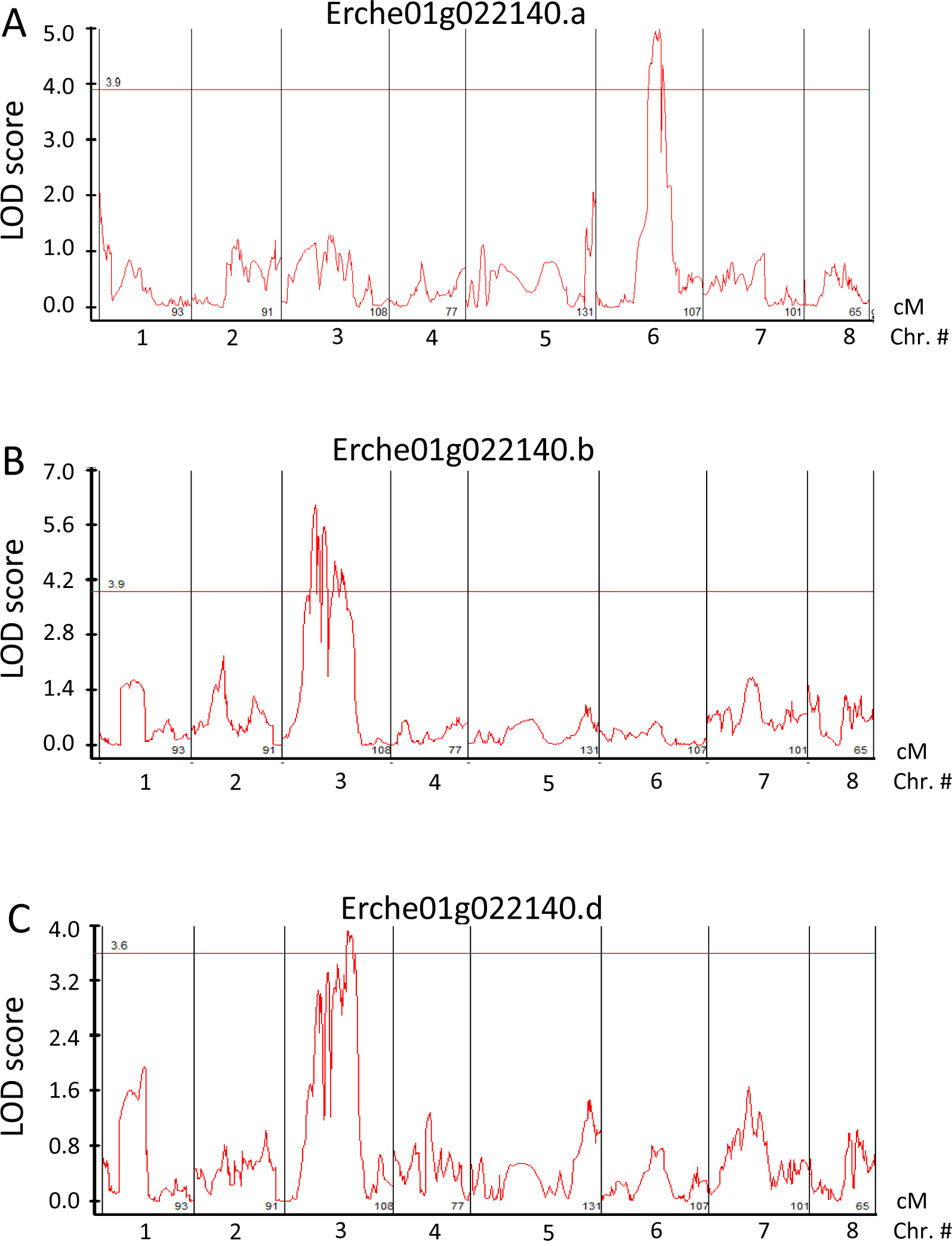
Quantitative trait loci (QTL) affecting indole glucosinolate methyltransferase gene expression in *Erysimum cheiranthoides*. LOD plots of (A) Erche01g022140.a, (B) Erche01g022140.b, and (C) Erche01g022140.c expression levels in an Elbtalaue x Konstanz F2 population are shown. Horizontal lines in are 95% confidence levels, calculated based on 500 permutations of the data. Graphs were created using Windows QTL Cartographer.

